# PRDX6 Modulates Immune Checkpoint Inhibitor Response by Antagonizing Ferroptosis Induced By HDAC Inhibitors

**DOI:** 10.64898/2026.01.21.700636

**Authors:** Zeyu Liu, Jessy John, Nickolas Johnson, Priya Singh, Ziyi Wang, Gilson J. Sanchez, Dylan Simons, Jing H Wang, Xuedong Liu

**Author notes:** These authors contributed equally (Zeyu Liu, Jessy John). Correspondence and Lead Contact (X.L.) and (J.W).

## Abstract

Therapeutic resistance limits the efficacy of histone deacetylase (HDAC) inhibitors and immune checkpoint therapies in cancer. While HDAC inhibitors can induce ferroptosis, tumor cells often evade this cell death via antioxidant defenses. Here we identify peroxiredoxin 6 (PRDX6) as a critical modulator of resistance to HDAC inhibitor largazole by suppressing ferroptosis through its phospholipase A2 activity and maintaining GPX4 expression. Using genome-wide CRISPR activation screening, biochemical assays, and syngeneic tumor models, we show that PRDX6 depletion enhances largazole-induced lipid peroxidation, ferroptotic stress, and reshapes the tumor microenvironment to promote T-cell infiltration and inflammatory cytokine release. Importantly, combining PRDX6 knockdown with HDAC inhibition potentiates anti-PD-L1 immunotherapy efficacy and prolongs survival *in vivo*. These findings reveal PRDX6 as a redox gatekeeper linking ferroptosis resistance to immune evasion and suggest that co-targeting PRDX6 and HDAC pathways may improve responses to cancer immunotherapy.

## INTRODUCTION

Despite remarkable advances in targeted therapies and immunotherapy, therapeutic resistance remains one of the most significant challenges in the effective treatment of solid tumors ^1–3^. A major barrier to achieving durable clinical responses is the ability of tumor cells to evade immune surveillance and adapt to therapeutic stress through epigenetic reprogramming and redox remodeling ^4–6^. Histone deacetylase (HDAC) inhibitors such as largazole or its derivatives, hold promise for reprogramming gene expression ^7–9^. These agents can reactivate silenced tumor suppressor pathways and enhance antitumor immunity^10,11^. However, their efficacy is often limited by adaptive resistance mechanisms that remain incompletely understood ^12–14^.

Recent studies have highlighted ferroptosis, a regulated form, iron-dependent and immunogenic cell death driven by lipid peroxidation, as a therapeutically exploitable vulnerability in cancer ^15,16^. However, cancer cells often evade ferroptosis by activating robust antioxidant defense mechanisms ^17^. The specific molecular regulators that suppress ferroptotic signaling during HDAC inhibitor treatment remain incompletely understood. In particular, the roles of antioxidant enzymes in limiting ferroptosis and influencing the tumor immune microenvironment are not well defined ^18^.

Here, we identify peroxiredoxin 6 (PRDX6), a multifunctional antioxidant enzyme with both glutathione peroxidase (GPx) and phospholipase A2 (PLA2) activities ^19^, whose elevated expression correlates with aggressive tumor phenotypes and poor patient prognosis ^20–23^, as a key suppressor of ferroptosis and immune activation in squamous cell carcinoma. We demonstrate that elevated PRDX6 mediates adaptive resistance to HDAC inhibitors by suppressing largazole-induced ferroptosis, through a mechanism that is independent of largazole’s HDAC inhibitory activity. Conversely, PRDX6 deficiency enhances tumor cell sensitivity to ferroptosis induced by HDAC inhibitors. This deficiency also remodels the tumor microenvironment by increasing cytokine release and promoting T-cell infiltration and differentiation following HDAC inhibition. These findings reveal a previously unrecognized link between ferroptosis regulation and immune modulation, which has direct implications for overcoming resistance to immune checkpoint blockade

## RESULTS

### Genome-wide CRISPRa high-throughput screening identifies PRDX6 as a mediator of resistance to largazole-induced cell death in HCT116 cells

Largazole, a potent naturally occurring class I HDAC inhibitor, undergoes hydrolysis to release its active thiol form (Figure 1A) ^24,25^. While its epigenetic effects are well documented, the broader cellular responses it triggers and the mechanisms by which tumor cells adapt to resist its action remain incompletely understood ^26–28^. To systematically identify genes conferring resistance to largazole, we employed a genome-wide CRISPR activation (CRISPRa) high-throughput screen in the HCT116 colorectal cancer cell line (Figure 1B). HCT116 cells stably expressing CRISPR-dCas9-VP64-GFP were transduced with a pooled sgRNA library and split into three groups: two experimental groups and one baseline control group frozen at day 0 to represent the original sgRNA distribution. Experimental groups were treated with either vehicle (DMSO) or 100 nM largazole for 12 days, with fresh medium containing DMSO or largazole replenished every three days. Following treatment, genomic DNA was extracted, and sgRNA abundance was quantified by next-generation sequencing. Data were analyzed using PinAPL-Py ^29^ to determine gene-level enrichment by comparing sgRNA frequencies between largazole- and DMSO-treated cells. Applying stringent statistical criteria (p-value < 2 × 10⁻L), we identified 202 genes significantly enriched under largazole selection (Figure 1C). Notably, ASS1 emerged as the top hit, consistent with previous studies showing that ASS1 depletion sensitizes cells to HDAC inhibitors, thereby validating the fidelity of our screen ^30^. To further confirm our results, we performed a secondary validation screen by subcloning sgRNA fragments enriched in the primary screen into the original CRISPRa vector to generate a focused sub-library. This sub-library was re-screened under the same conditions, and sgRNA enrichment was again assessed by comparing results to the primary screen. The strong and reproducible enrichment of an ASS1-specific sgRNA further confirmed the robustness of our screening approach (Figure 1D).

**Figure 1.**
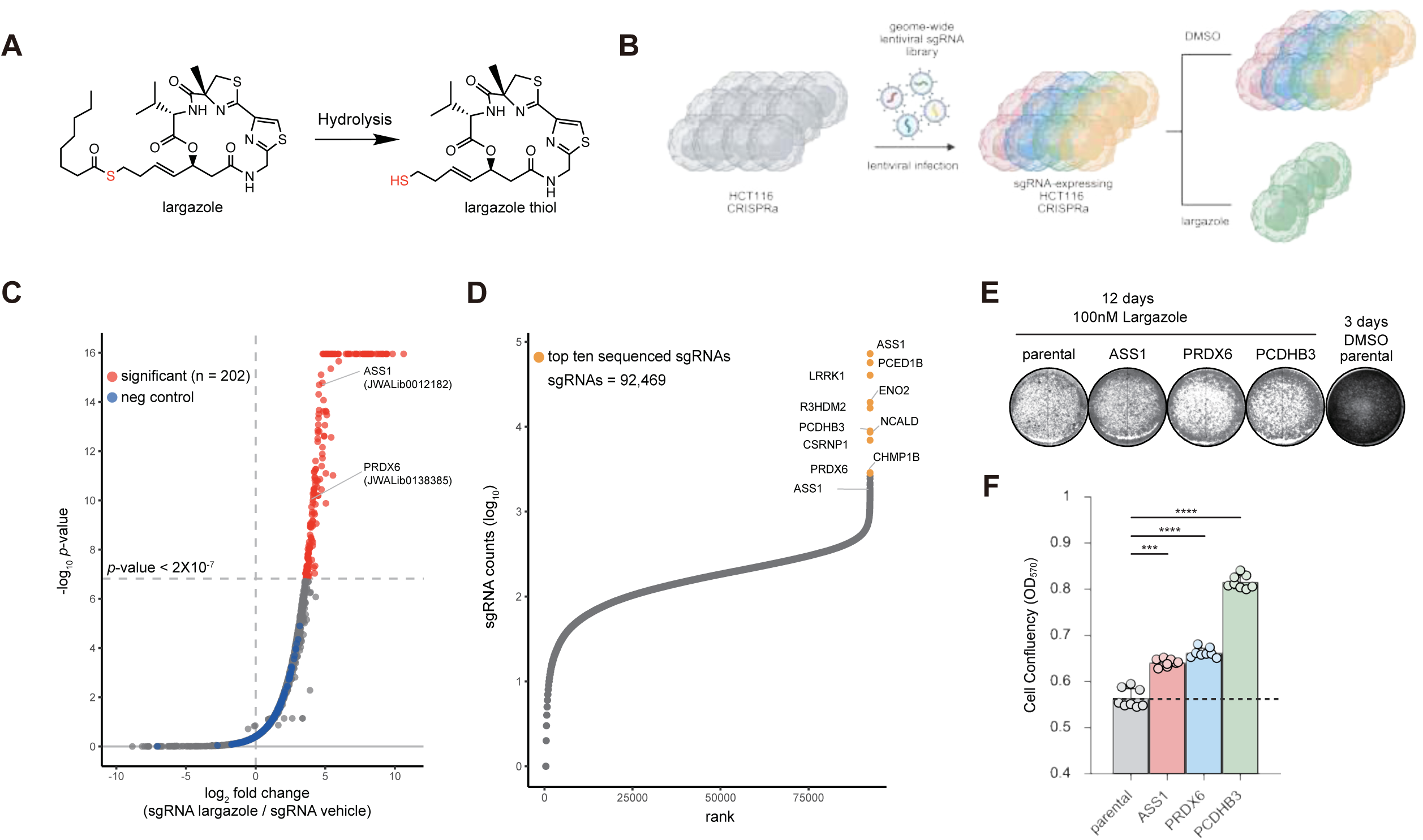
Genome wide CRISPRa high throughput screen identifies PRDX6 as a largazole target. **(A)** Structures of the prodrug largazole and its active form, largazole thiol, after hydrolysis. **(B)** Schematic of the genome wide lentiviral pooled screen for largazole. HCT116 cells expressing CRISPRa (dCas9-VP64-GFP) were infected with a lentiviral sgRNA library. After selection with 2 µg/mL puromycin, cells were split into two groups: one treated with DMSO and the other with 100 nM largazole. Following treatment, cells were lysed and genomic DNA was isolated, processed, and sequenced. **(C)** Differential sgRNA abundance analysis comparing largazole treated versus DMSO treated groups. A significance threshold of *p*-value < 2×10^-7^ was used to call hits. **(D)** Gene level enrichment from the secondary screen comparing DMSO and largazole conditions. Genes were ranked by the summed read counts of their sgRNAs. **(E)** Validation of screening results using cells overexpressing individual genes. HCT116 wild type cells with the indicated cDNA overexpression were treated with 100 nM largazole for 12 days, fixed with 4% formaldehyde in PBS, and stained with 0.05% crystal violet. A 3-day DMSO treatment of HCT116 wild type cells served as a natural growth control. Images were inverted to grayscale to enhance visibility. **(F)** Quantification of the crystal violet assays shown in panel E. Stain was solubilized and absorbance measured at 570 nm. Data are mean ± SD. Welch one way ANOVA with Games Howell multiple comparisons test was used. \*\*\**p* < 0.001, \*\*\*\**p* < 0.0001.

For individual validation, we selected 5 of the top 10 candidate genes from the secondary screen and generated stable HCT116 cell lines overexpressing each gene using a mammalian piggyback expression vector. These lines were treated with largazole under the same conditions as the screen, and cell viability was assessed based on final confluency (Figures 1E and 1F). Among the validated hits, overexpression of PRDX6 and PCDHB3 significantly increased resistance to largazole-induced cytotoxicity. We speculate that PCDHB3 may confer resistance by strengthening cell-cell adhesion or enhancing cytoskeletal integrity, which could potentially reduce susceptibility to apoptosis. However, further investigation is needed to confirm this mechanism. We focus on PRDX6 given its known role in regulating lipid peroxidation through its phospholipase A2 (PLA2) activity ^19^ and prior studies linking HDAC inhibition to ferroptosis and lipid peroxidation ^31^. We hypothesize that PRDX6 promotes resistance to largazole by mitigating ferroptosis through suppression of lipid peroxidation, a mechanism that we explore in detail in subsequent experiments.

### Overexpression of PRDX6 confers resistance to largazole without altering HDAC inhibition

To further evaluate the role of PRDX6 in mediating resistance to largazole, we generated HCT116 cell lines stably overexpressing either PRDX6 or ASS1, alongside an ASS1 knockdown (KD) line. Expression levels were confirmed by western blot analysis (Figure S1A). Largazole sensitivity assays revealed that both PRDX6- and ASS1-overexpressing cells exhibited increased resistance to largazole-induced cytotoxicity compared to wild-type controls, whereas ASS1 knockdown sensitized cells to largazole, consistent with previous findings ^30^ (Figures S1B and S1C).

To determine whether the observed resistance was due to altered HDAC inhibition, we assessed histone acetylation levels following a 2-hour treatment with increasing concentrations of largazole. Specifically, we examined pan-H4ac (a marker of global histone acetylation), H3K9ac (linked to promoter activity), and H3K27ac (associated with enhancer activation). Western blot analysis revealed minor differences in histone acetylation patterns at the lowest largazole concentration tested (5 nM) in cells overexpressing ASS1 or PRDX6 compared to wild-type cells. However, these subtle differences were no longer observable at higher largazole concentrations (Figure S1D). Given that our original CRISPRa screen utilized 100 nM largazole, a concentration substantially higher than the threshold (∼10 nM) at which the acetylation differences dissipate, we conclude that the resistance to largazole conferred by ASS1 and PRDX6 overexpression is independent of any significant alterations in HDAC inhibitory activity. Thus, PRDX6-mediated resistance likely involves additional mechanisms, separate from the canonical HDAC inhibition pathway.

### PLA2 function of PRDX6 modulates cellular sensitivity to largazole

Given that ASS1 KD sensitized HCT116 cells to largazole, we next investigated if PRDX6 KD produces a similar sensitization effect and whether this sensitivity depends on the enzymatic functions of PRDX6. To address this, we generated stable PRDX6 KD HCT116 cells using shRNA and performed rescue experiments with cDNAs immune to shRNA encoding either wild-type PRDX6 (PRDX6-WT), a PLA2-inactive mutant (S32A), a GPx-inactive mutant (C47S), or a double mutant (S32A/C47S) lacking both catalytic activities ^19^. Largazole sensitivity was assessed using the same treatment protocol as our original CRISPR screen, with cell viability measured by final confluency measurements (Figures 2A and 2B). Consistent with ASS1 KD, PRDX6 KD significantly increased sensitivity to largazole, which could be partially rescued by re-expression of PRDX6-WT. Notably, the PLA2-inactive S32A mutant showed impaired rescue compared to PRDX6-WT, whereas GPx-inactive C47S mutant restored resistance to a similar degree as PRDX6-WT. The double mutant (S32A/C47S) failed to rescue largazole sensitivity altogether. The greater loss of rescue in the double mutant compared to the S32A mutant alone may reflect additional structural or functional disruption beyond PLA2 inactivation. In contrast, the C47S mutant retains PLA2 activity and overall protein integrity, enabling effective rescue. However, further study is needed to confirm this mechanism. As these results were observed in the absence of any significant changes in HDAC inhibition across cell lines (Figure 2C), we conclude that PRDX6 confers resistance to largazole primarily through its PLA2 activity, independent of GPx function or direct modulation of HDAC activity.

**Figure 2.**
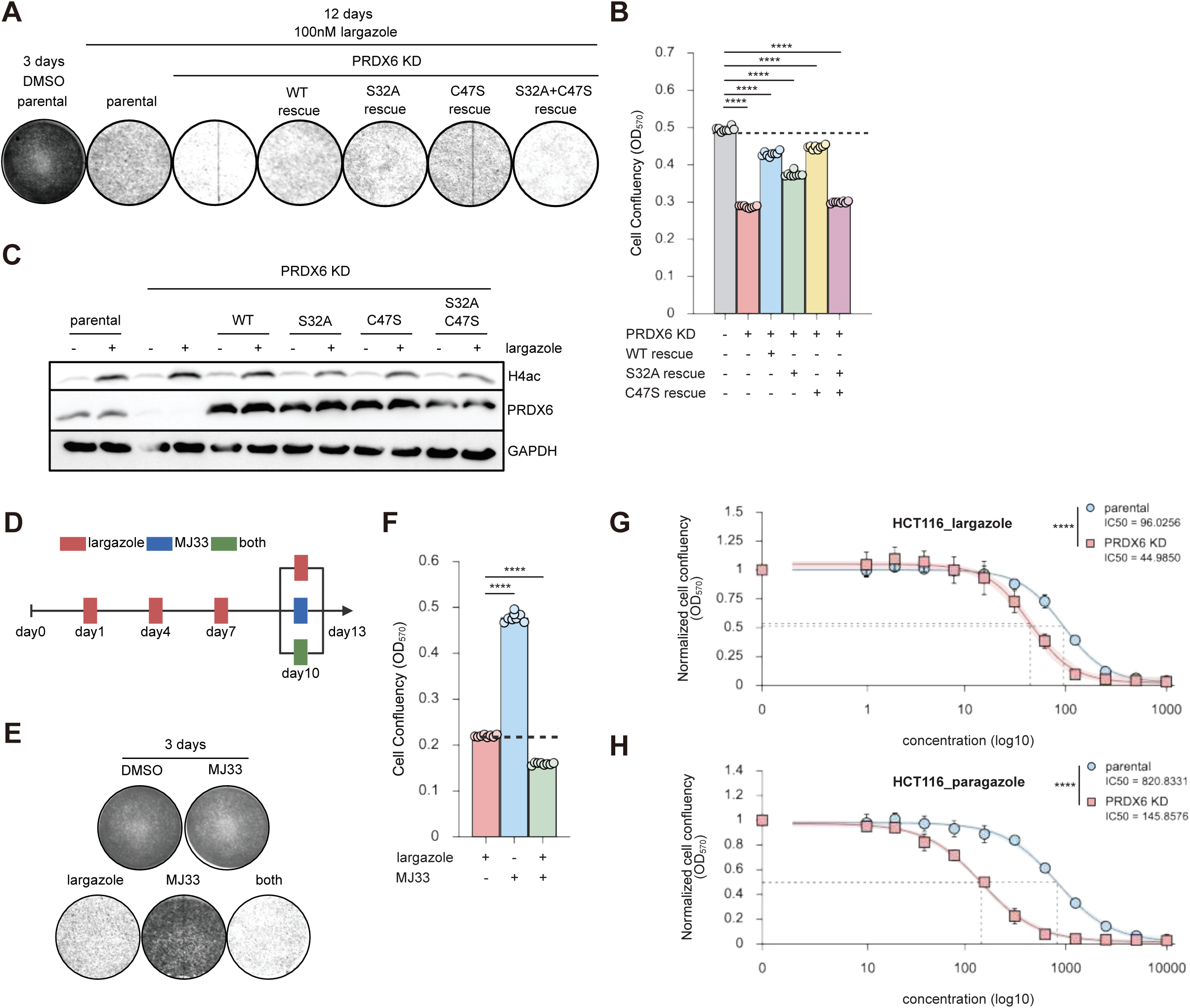
PRDX6 alters cell sensitivity to largazole without comprising HDAC inhibition. **(A)** Largazole sensitivity assay in cells with PRDX6 knockdown and the indicated rescues. HCT116 cells with PRDX6 knockdown and PRDX6 knockdown rescued with WT, S32A, C47S, or S32A+C47S were treated with 100 nM largazole for 12 days, fixed with 4% formaldehyde in PBS, and stained with 0.05% crystal violet. A 3-day DMSO treatment of HCT116 parental cells served as a natural growth control. Images were inverted to grayscale to enhance visibility. **(B)** Quantification of panel A. Crystal violet stain was solubilized and absorbance measured at 570 nm. Data are mean ± SD. One way ANOVA with Tukey multiple comparisons test was used. \*\*\*\**p* < 0.0001. **(C)** Western blot of pan H4ac levels in HCT116 parental, PRDX6 knockdown, and PRDX6 knockdown cells with the indicated rescues. Cells were treated with 100 nM largazole for 2 hours. GAPDH was used as a loading control. **(D)** Schematic of the combination treatment assay. 3×10^5^ HCT116 parental cells were seeded per well in a 6 well plate. All wells were treated with 100 nM largazole for 9 days, then washed with PBS and replaced with 100 nM largazole, 10 µM MJ33, or 100 nM largazole plus 10 µM MJ33 for 3 additional days. **(E-F)** Results and quantification of panel D. 3-day DMSO and MJ33 treatments of HCT116 parental cells were used to indicate natural growth rate and any growth effect of MJ33 alone. Images were inverted to grayscale to enhance visibility. Crystal violet stain was solubilized and absorbance measured at 570 nm. Data are mean ± SD. One way ANOVA with Tukey multiple comparisons test was used. \*\*\*\**p* < 0.0001. **(G-H)** IC_50_ values for largazole and paragazole in HCT116 parental versus PRDX6 knockdown cells. HCT116 parental and PRDX6 knockdown cells were seeded into a 96 well plate in 4 lanes × 12 wells with 4,000 cells per well. Twelve hours after seeding, medium was replaced with fresh DMEM containing a serial dilution of largazole, and cells were treated for 3 days. Cells were then washed with PBS, fixed with 4% formaldehyde in PBS, stained with 0.05% crystal violet, solubilized, and absorbance was read at 530 nm. Wells without largazole were used as controls for normalization. Data are mean ± SD. A four-parameter logistic regression was used to calculate IC_50_. Student *t*-test was used for statistical analysis. \*\*\*\**p*<0.0001.

To further confirm that PRDX6’s PLA2 activity is responsible for modulating largazole sensitivity, we performed a combination treatment of largazole with MJ33 ^32^, a reversible inhibitor of PRDX6 PLA2 activity, to phenocopy the effect observed with the S32A mutation ^33,34^. Briefly, HCT116 WT cells were treated with 100 nM largazole for 9 days, followed by 3 days of exposure to either 100 nM largazole alone, 10 µM MJ33 alone, or the combination of both agents (Figure 2D). After this treatment period, cells receiving the combination of largazole and MJ33 exhibited significantly reduced viability, confirming that inhibition of PRDX6’s PLA2 function enhances largazole-induced cytotoxicity (Figures 2E and 2F). In the short-term growth inhibition assay, PRDX6 knockdown also sensitizes cells to both largazole and an analog of largazole known as paragazole/OKI-5 (Figure 2G and 2H).

### PDRX6 suppresses largazole-induced lipid peroxidation and ferroptosis

The PLA2 activity of PRDX6 is known to reduce peroxidized lipid accumulation by removing oxidized sn-2 fatty acyl chains from phospholipids, thereby facilitating membrane repair and aiding redox homeostasis ^19,35^. Since increased lipid peroxidation can induce ferroptosis ^15^, we investigated whether largazole triggers ferroptotic signaling and if this response is enhanced in the absence of PRDX6. To address this, we first quantified lipid peroxidation in HCT116 WT and PRDX6 KD cells using an imaging-based assay with the Image-iT® Lipid Peroxidation Sensor. Cells were treated with DMSO, cumene hydroperoxide (CHP, a positive control), or largazole for defined time intervals, followed by fixation and staining with Hoechst 33342. The lipid peroxidation sensor shifts fluorescence from TRITC to FITC upon oxidation, allowing ratiometric measurement of peroxidation levels via FITC/TRITC signal. As expected, CHP treatment (1 hour) elevated lipid peroxidation in both WT and PRDX6 KD cells, but levels were significantly higher in PRDX6-deficient cells, confirming increased oxidative vulnerability due to PRDX6 loss (Figures 3A and 3B). When cells were treated with largazole, lipid peroxidation progressively increased in both WT and PRDX6 KD cells at 18 hours post-treatment, but this response was again more pronounced in PRDX6 KD cells, supporting the hypothesis that largazole induces lipid peroxidation and PRDX6 protects against largazole-induced lipid oxidative stress (Figures 3A and 3B).

**Figure 3.**
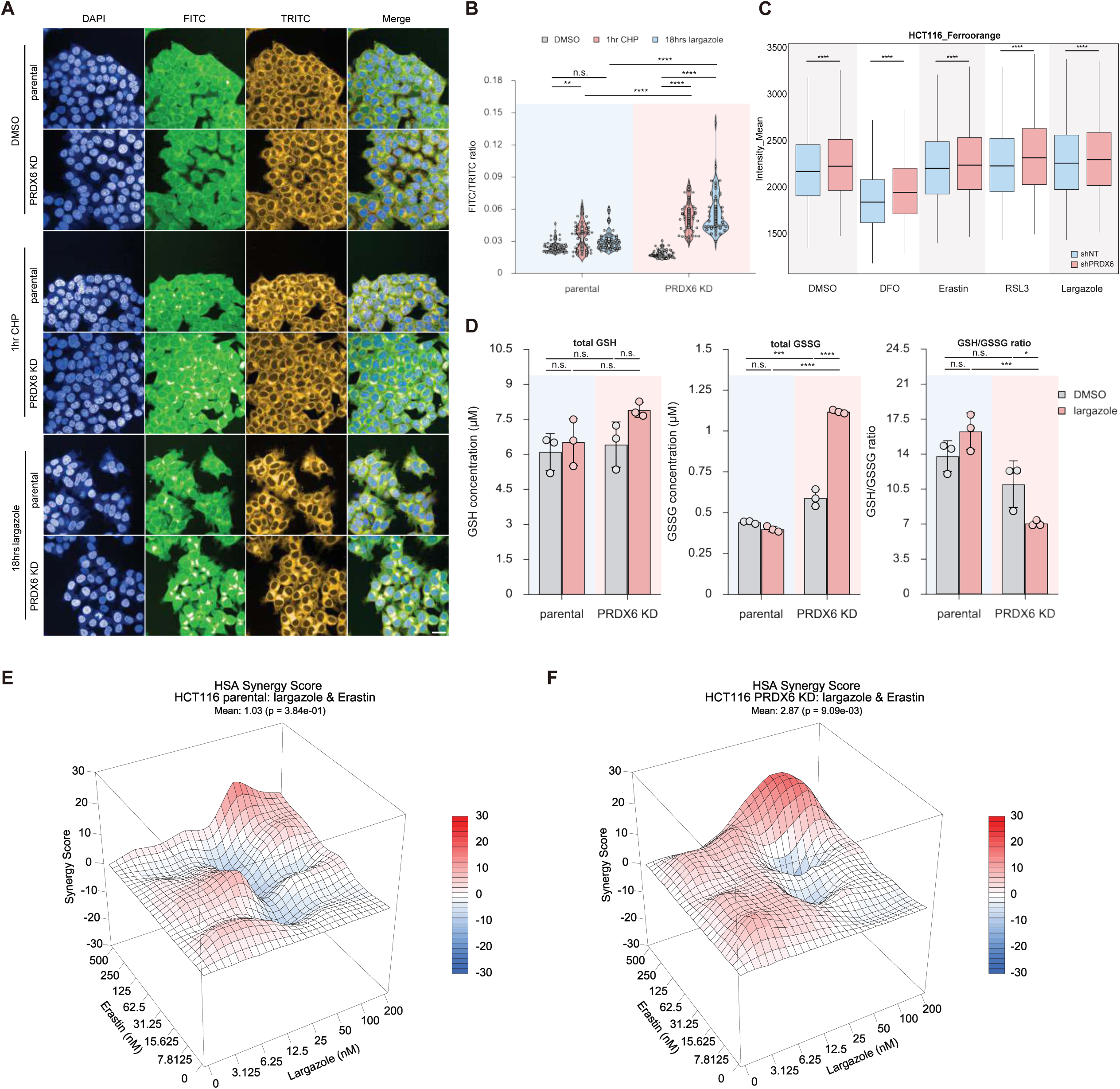
PRDX6 knockdown increases vulnerability to largazole induced lipid peroxidation stress. **(A)** Largazole induces lipid peroxidation in HCT116 cells. HCT116 parental and PRDX6 knockdown cells were seeded in a 96 well plate at 10,000 cells per well. After 24 hours, medium was replaced with fresh DMEM containing the indicated treatments for the indicated durations. Cells were then washed three times with PBS, fixed with 4% paraformaldehyde in PBS, and stained with 10 µM Image iT Lipid Peroxidation Sensor and 10 µg/mL Hoechst 33342 for 3 days at 4°C in the dark. After incubation, cells were washed three times with PBS and imaged using DAPI, FITC, and TRITC channels. Scale bar, 100 µm. **(B)** Quantification of panel A. Fifty cells per condition were randomly selected and the mean FITC and TRITC fluorescence intensities per cell were measured. The FITC/TRITC ratio was used as a readout of lipid peroxidation stress. Two-way ANOVA with Tukey multiple comparisons test was used. \*\**p* < 0.01, \*\*\*\**p* < 0.0001, n.s.= not significant. **(C)** PRDX6 knockdown induces iron accumulation in HCT116 cells upon largazole treatments. After indicated treatments, live cells were stained with 1LµM FerroOrange (Dojindo) and costained with 10Lµg/mL Hoechst 33342 for 30 minutes prior to imaging, without washout. Imaging was performed on the Revvity Opera Phenix Confocal System using a 20X water objective and standard settings. Nuclei were segmented using Hoechst signal, and perinuclear cytoplasmic regions were defined to quantify FerroOrange fluorescence on a single-cell basis. Border cells were excluded, and integrated fluorescence intensity per cell was calculated. “High” Fe²⁺ cells were defined using Otsu’s threshold from control wells, and the percentage of high-signal cells was quantified per well. Bars represent mean ± SEM from biological replicates. Statistical significance was determined by two-way ANOVA with Tukey’s post-hoc test. \*\*\**p* < 0.001, \*\*\*\**p* < 0.0001, n.s.= not significant. **(D)** Largazole increases GSSG in PRDX6 knockdown HCT116 cells. HCT116 parental and PRDX6 knockdown cells were seeded in a 96 well plate at 10,000 cells per well. After 24 hours, medium was replaced with fresh DMEM containing 100 nM largazole for 18 hours. GSH and GSSG were measured using the GSH/GSSG Glo assay, with readings normalized using the CellTiter Glo luminescent viability assay. Two way ANOVA with Tukey multiple comparisons test was used. \**p* < 0.05, \*\*\**p* < 0.001, \*\*\*\**p* < 0.0001, n.s. = not significant. (**E-F**) Largazole shows increased synergy with erastin in PRDX6 knockdown HCT116 cells. HCT116 parental and PRDX6 knockdown cells were seeded in a 96 well plate in 8 lanes × 8 wells at 8,000 cells per well. Twelve hours after seeding, medium was replaced with fresh DMEM containing serial dilutions of largazole and erastin2, and cells were treated for 3 days. Cells were then washed twice with PBS, fixed with 4% formaldehyde in PBS, stained with 0.05% crystal violet, solubilized, and absorbance was read at 530 nm. Wells without treatment served as normalization controls. Synergy scores were computed and visualized using SynergyFinder+.

We next asked whether largazole-induced oxidative stress is linked to ferroptosis by evaluating cytosolic mobilized iron accumulation, a hallmark of ferroptosis ^15,17^. Largazole treatment induced iron influx in both WT and PRDX6 KD cells, with a more robust response in PRDX6 KD cells (Figures 3C). Another hallmark of ferroptosis is the depletion of intracellular glutathione (GSH) ^15,17^. We measured the intracellular redox status via the glutathione/glutathione disulfide (GSH/GSSG) ratio. PRDX6 KD alone induced only modest increases in GSSG, with minimal impact on the overall GSH/GSSG balance (Figure 3D). However, after 18 hours of largazole exposure, PRDX6-deficient cells exhibited a marked elevation in GSSG and a significant drop in the GSH/GSSG ratio, further indicating enhanced ferroptotic stress under these conditions (Figure 3D).

To determine whether PRDX6 deficiency enhances susceptibility to ferroptosis in general, we treated WT and PRDX6 KD cells with well-established ferroptosis inducers, erastin2 and RSL3 ^36,37^, in combination with serial dilutions of largazole. PRDX6 KD alone significantly increased sensitivity to erastin2, whereas no difference was observed for RSL3 (Supplemental Figures S2A-S2D). However, when combined with largazole, the cytotoxic effects of both inducers were further potentiated, with HSA synergy scores elevated in PRDX6 KD cells across defined dose ranges (Figure 3E and 3F; Supplemental Figures S2E and S2F).Together, these results demonstrate that PRDX6 suppresses largazole-induced ferroptosis by limiting lipid peroxidation, iron accumulation and GSSG depletion, which are three hallmarks of ferroptosis. Its depletion creates a ferroptosis-permissive state that sensitizes tumor cells to HDAC inhibition and opens the door to combination strategies with ferroptosis inducers.

### PRDX6 deficiency increases cell vulnerability to ferroptosis by lowering GPX4

Building on our findings that PRDX6 modulates largazole sensitivity through its regulation of lipid peroxidation, we next investigated the broader clinical significance of PRDX6 expression across tumor types. Analysis of The Cancer Genome Atlas (TCGA) dataset revealed that high PRDX6 expression is significantly associated with reduced overall survival across cancers (Figure 4A), implicating it as a potential driver of disease progression. Stratifying by tumor type, we identified seven cancers where elevated PRDX6 expression correlated strongly with poor prognosis, with head and neck squamous cell carcinoma (HNSCC) showing the most pronounced dependency (Figure 4B; Supplemental Figure S3A and S3B). In contrast, other peroxiredoxin family members showed limited prognostic relevance in selected cancers, reinforcing the unique clinical importance of PRDX6 (Supplemental S4).

**Figure 4.**
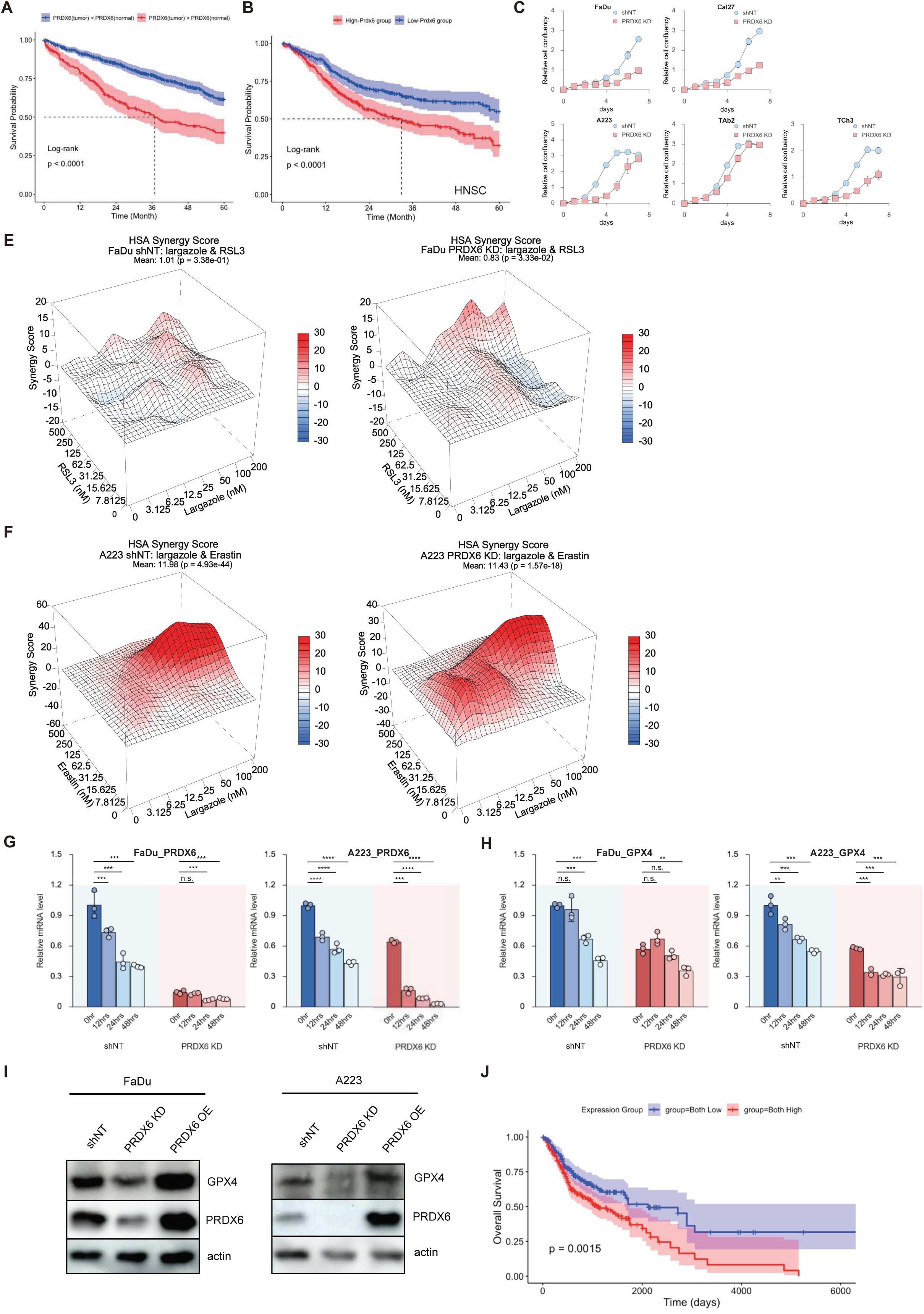
PRDX6 is critical to maintain GPX4 level in head and neck squamous cell carcinoma. **(A)** Overall pan-cancer survival analysis stratified by PRDX6 expression. Patients were grouped by significantly upregulated versus significantly downregulated PRDX6 in tumor relative to matched normal tissue. Shaded bands indicate 95% confidence intervals. Log-rank test *p*-value < 0.05. **(B)** PRDX6 expression level is critical for survival probability for patients with head and neck squamous cell carcinoma (HNSCC). Five year-Kaplan-Meier overall survival from TCGA data stratified by median PRDX6 expression (High PRDX6, red; Low PRDX6, blue). Shaded bands represent 95% confidence intervals. Log-rank test *p*-value < 0.05. **(C-D)** PRDX6 knockdown slows proliferation in squamous carcinoma cell lines. For all indicated cell lines, 12 wells of 500 cells were seeded into 7 plates of 96-well plate (1 plate/day). After indicated duration of incubation, plates were fixed and crystal violet staining was performed. The plates were solubilized, and absorbance was measured at 570 nm. Data are mean± SD. **(E-F)** Largazole shows synergistic effect with RSL3 and with erastin2 in A223 and FaDu PRDX6 knockdown cells. For both FaDu and A223, shNT and PRDX6 knockdown cells were seeded into 8 lanes x 8 wells of 96-well plate with 8,000 cells in each well. 12 hours after seeding, DMEM was replaced by fresh DMEM with largazole and RSL3 or and erastin2 in serial-diluted manner, and cells were treated for 3 days. After 3 days treatment, cells were washed with PBS twice, fixed with 4% formaldehyde in PBS, stained with 0.05% crystal violet, solubilized and quantified by reading absorbance for each well at 570 nm. Wells with no treatment were used as control to normalize the data and synergy scores were calculated and visualized using SynergyFinder+. **(G-H)** Largazole downregulates PRDX6 and GPX4 mRNA in FaDu and A223 cells. FaDu and A223 cells with either shNT or PRDX6 knockdown were treated with 100LnM largazole for the indicated times. Total RNA was extracted at each time point and subjected to RT-qPCR to assess PRDX6 and GPX4 mRNA expression. β-actin was used as the housekeeping gene for normalization. mRNA levels were normalized to the respective 0Lhr WT control group for both cell types. Data represent the relative expression of PRDX6 and GPX4 over time in response to largazole treatment. A regular one-way ANOVA was performed for statistical analysis. \*\**p* < 0.01, \*\*\**p* < 0.001, \*\*\*\**p* < 0.0001, n.s. = not significant. **(I)** PRDX6 regulates GPX4 protein level. Western blot for GPX4 expression levels in A223 or FaDu WT, PRDX6 KD and PRDX6 overexpression (OE) cell lines. Actin was used as a loading control. **(J)** PRDX6 and GPX4 co-expression and survival in HNSCC. Five year Kaplan Meier overall survival from TCGA stratified by combined PRDX6 and GPX4 expression (Both high, red; Both low, blue) using median cutoffs. Shaded bands indicate 95 percent confidence intervals. Log rank test *p* < 0.05.

To functionally validate these observations and elucidate a potential biochemical mechanism for poor prognosis associated with elevated PRDX6 expression, we knocked down PRDX6 in two representative HNSCC lines (FaDu and CAL-27) and three mouse SCC lines (A223, TAb2, and TCh3). In all cases, we observed a marked suppression of cell proliferation (Figure 4C and 4D). Consistent with our earlier findings, PRDX6 KD sensitized all five lines to largazole (Table 1). Moreover, depletion of PRDX6 in HNSCC cell lines potentiates ferroptotic responses as observed in colorectal models: increased lipid peroxidation, elevated iron accumulation, and heightened synergy with ferroptosis inducers, all consistent with enhanced ferroptotic vulnerability (Figures 4E and 4F; Supplemental Figures S5 and S6). These data affirm PRDX6 as a ferroptosis suppressor whose loss unleashes a significant vulnerability to HDAC inhibition in HNSCC.

To further dissect the molecular basis of this vulnerability, we turned to the DepMap CRISPR dependency dataset ^38^. While PRDX6 displayed only mild essentiality across most HNSCC lines, some established cancer cell lines such as LB771HNC, exhibit strong PRDX6 dependence (Supplemental Figures S7A–B). Strikingly, GPX4, one of the most well-established negative regulators of ferroptosis, ranked among the top 10 essential genes in this same cell line (Supplemental Figure S7C; Table 2). When we plotted PRDX6 and GPX4 dependency scores across HNSCC lines, we found a clear co-dependency pattern: cell lines such as FaDu and CAL-27 showed mutual reliance on both genes (Supplemental Figure S7D), pointing to a potential functional relationship. Co-expression analysis at the mRNA level further supported this link, with PRDX6 and GPX4 expression positively correlated in HNSCC tumors (Supplemental Figures S7E–H).

To delineate molecular basis of PRDX6 and GPX4 interdependence, we measured PRDX6 and GPX4 mRNA levels in WT and PRDX6 KD FaDu and A223 cells following largazole treatment over time. Consistent with our co-expression analysis, largazole repressed the expression of both genes, and a low basal level of PRDX6 further amplified this effect (Figure 4G and 4H). Western blots further confirmed that protein expression levels of PRDX6 and GPX4 are directly connected. Depletion of PRDX6 reduced GPX4 protein expression, whereas overexpression of PRDX6 increased GPX4 protein levels in both human FaDu and mouse A223 HNSCC lines (Figure 4I). The coordinated protein expression could in part be attributed to the reported selenium carrier function of PRDX6 in promoting GPX4 translation ^39–41^. Our co-expression analysis also points to a significant new mechanism of co-regulation at the level of mRNA expression upstream of protein translation.

To further define the clinical and translational significance of PRDX6 and GPX4 co-regulation, we analyzed TCGA survival data and found that patients with low expression of both PRDX6 and GPX4 had significantly improved overall survival, supporting a combined role for these genes in disease progression and therapy resistance (Figure 4J). We also examined the relationship between PRDX6 and GPX4 in patient samples from a phase II clinical trial that evaluated entinostat plus nivolumab in advanced pancreatic ductal adenocarcinoma ^42^. Like largazole and its derivative paragaozle/OKI-5, entinostat is a class I selective HDAC inhibitor. RNA-seq data were available for a subset of patients before and after treatment (Supplemental Figure S8A) ^42^. Although the number of patient samples was limited, there was a trend toward reduced PRDX6 and GPX4 mRNA after treatment (Supplemental Figure S8B). Notably, the three patients who responded to therapy showed the greatest reduction in both PRDX6 and GPX4 (patients 18 and 38; RNA for patient 28 was not available) (Supplemental Figure S8B) ^42^. We plotted PRDX6 and GPX4 mRNA levels before and after treatment, together with two other PRDX family members. PRDX6 and GPX4 showed a significant correlation at baseline (p < 0.02), which became more significant after entinostat and nivolumab treatment (p < 0.004) (Supplemental Figure S8C and S8D). No significant correlation was observed between GPX4 and PRDX2 or PRDX4. Collectively, these results support a mechanistic axis in which PRDX6 maintains GPX4 mRNA and protein expression and redox homeostasis, thereby buffering cells from ferroptotic stress. The coordinated expression of PRDX6 and GPX4 may be susceptible to targeting by HDAC inhibition *in vitro* and *in vivo*.

### PRDX6 deficiency primes tumor cells for ferroptosis and promotes inflammatory responses upon largazole treatment

To gain further mechanistic insights, we performed RNA-sequencing (RNA-seq) analysis on A223 parental and PRDX6 KD cells treated with DMSO or 100 nM largazole for 18 hours. Pathway enrichment analyses indicated that both largazole treatment and PRDX6 depletion activate overlapping inflammatory signaling pathways, including interferon gamma (IFN-γ) and interferon alpha (IFN-α) pathways, with additive effects observed upon combined PRDX6 KD and largazole treatment (Figure 5A). Differential expression analysis further revealed elevated expression of the whole inflammatory pathway and a complete turnover of ferroptosis pathway activation in PRDX6 KD cells with largazole treatment (Figure 5B and 5C). Basically, largazole treatment to PRDX6 KD cells induces a ferroptosis-prone state with robust activation of innate immune and inflammatory signaling pathways. Suppressed antioxidant and lipid-protective mechanisms, combined with increased labile iron, sensitizes cells to lipid peroxidation and oxidative stress, creating a permissive environment for ferroptotic cell death. Concurrently, elevated expression of cytokines, chemokines, interferon-stimulated genes, and pattern recognition receptors reflects heightened innate immune surveillance and stress response, including inflammasome activation and type I interferon signaling. This transcriptional program suggests that ferroptosis sensitization occurs in concert with innate inflammatory responses, positioning the treated cells at the intersection of regulated cell death and immunogenic signaling.

**Figure 5.**
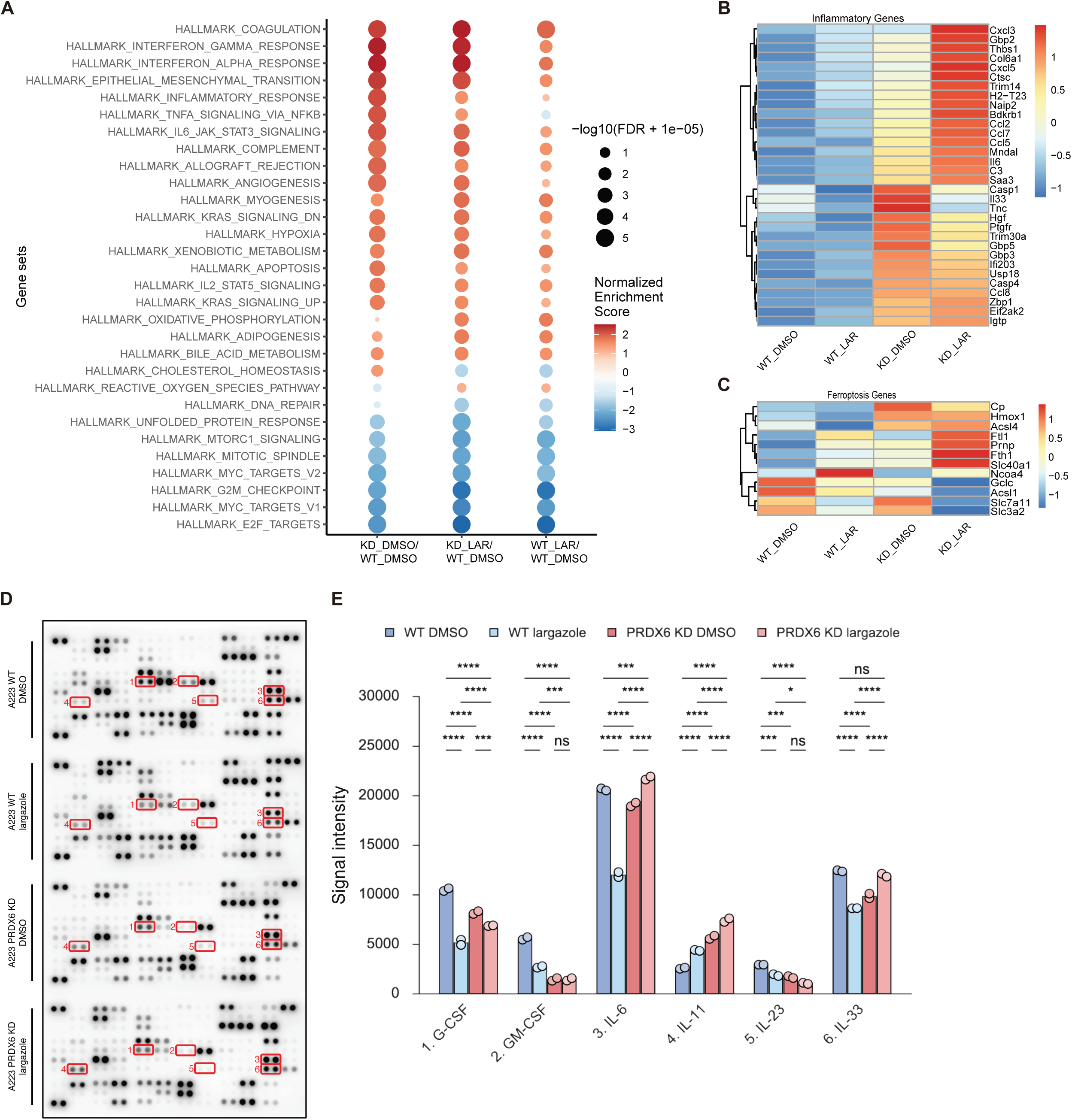
PRDX6 restrains largazole induced inflammatory and ferroptotic responses in squamous carcinoma cells. **(A)** GSEA shows that PRDX6 knockdown in A223 cells enhances largazole induced inflammatory responses. Bubble plot summarizing GSEA results of Hallmark gene sets enrichment across three conditions: KD_DMSO vs WT_DMSO, WT_LAR vs WT_DMSO and KD_LAR vs KD_DMSO. Each point represents a significantly enriched pathway (FDR < 0.25). Color indicates the normalized enrichment score (NES), reflecting the direction and magnitude of enrichment. Bubble size reflects the negative log10 of the adjusted *p*-value. **(B-C)** Largazole induces inflammatory signaling and ferroptosis in a PRDX6 knockdown dependent manner. Heatmaps display expression of genes in inflammation and ferroptosis pathways across the indicated conditions. Gene values were taken from DESeq2 normalized counts, log2 transformed, and row scaled (z score). The color scale represents relative expression (blue, low; red, high). Only genes detected in the RNA seq dataset and present in the normalized matrix are shown. **(D)** Cytokine array reveals elevated inflammatory responses induced by largazole treatment in A223 PRDX6 KD cells. A223 WT and PRDX6 KD cells were seeded into a 6-well plate with 2×10^6^ cells in each well. After 24 hours, medium was replaced with 1 mL fresh DMEM containing DMSO or 100 nM largazole for 18 hours. Cells were collected and cytokine levels were measured using the Proteome Profiler Array Mouse XL Cytokine kit. **(E)** Quantification of panel D. Spot intensities were measured and analyzed by two way ANOVA with Tukey multiple comparisons. \**p* < 0.05, \*\*\**p* < 0.001, \*\*\*\**p* < 0.0001, n.s. = not significant.

To validate these RNA-seq findings and explore cytokine-mediated immune modulation, we performed cytokine array analyses using lysates from A223 WT and PRDX6 KD cells treated with DMSO or largazole. Confirming our transcriptomic data, we observed significant modulation of IL-6 and IL-33 (Figure 5D, 5E3, and 5E6). Additionally, an increase in IL-23 and decrease in IL-11 were observed, suggesting a potential adaptive resistance mechanism against largazole (Figure 5E4 and 5E5). Moreover, largazole treatment in PRDX6 KD cells induced pronounced alterations in tumor microenvironment associated cytokines, including decreased levels of immunosuppressive cytokines G-CSF, GM-CSF, M-CSF, GDF-15, and osteopontin, alongside increased pro-inflammatory chemokines CCL17 and CXCL2 (Figure 5E1, 5E2, and Supplemental Figure S9A and S9B). These changes indicate enhanced potential for T-cell infiltration and reduced myeloid cells-driven immunosuppression. Collectively, our results suggest that combining PRDX6 KD with largazole treatment reprograms the tumor-intrinsic stress response toward a ferroptosis-permissive, immunostimulatory state, marked by heightened inflammatory gene expression, altered cytokine secretion, and reduced immunosuppressive signaling.

### PRDX6 depletion sensitizes SCC tumor growth inhibition by HDAC inhibitor OKI-179 and prolongs host survival *in vivo*

To assess whether PRDX6 KD sensitize tumor growth inhibition by HDAC inhibitor *in vivo*, we utilized a previously characterized syngeneic transplantable mouse tumor model (A223), which was derived from a Smad4 deficiency squamous tumor and has been shown to be a suitable animal model for HNSCC ^43–45^. For *in vivo* studies, we employed OKI-179, an orally bioavailable analog derived from paragazole/OKI-5 that is optimized for therapeutic applications and being evaluated in phase I/II clinical trials (NCT05340621) ^28^. Briefly, A223 parental or A223 PRDX6 KD tumor cells were implanted in the flank of WT C57BL/6 (B6) mice. WT B6 mice, inoculated with A223 or A223 PRDX6 KD tumors, received 60 mg/kg OKI-179 every two days once the tumor size reached ∼180 mm^3^. In untreated groups (Figure 6A and 6B), PRDX6 KD tumors were consistently growing slower than wild type counterparts (dark blue vs dark red) though not statistically significant, indicating PRDX6 depletion slows tumor growth. While WT A223 tumors were sensitive to OKI-179 treatment (Figure 6A, dark blue vs light blue), A223 PRDX6 KD tumors were significantly more sensitive (Figure 6A, dark red vs light red and light blue vs light red). On Day 46, when the study was concluded, mice with parental A223 tumors showed no survival difference between the two groups with or without OKI-179 treatment. In contrast, most mice harboring A223 PRDX6 KD tumors treated with OKI-179 were still alive (>80%), whereas the untreated group exhibited much lower survival (Figure 6C). These results suggest that PRDX6 depletion enhances tumor growth meditated by OKI-179.

**Figure 6.**
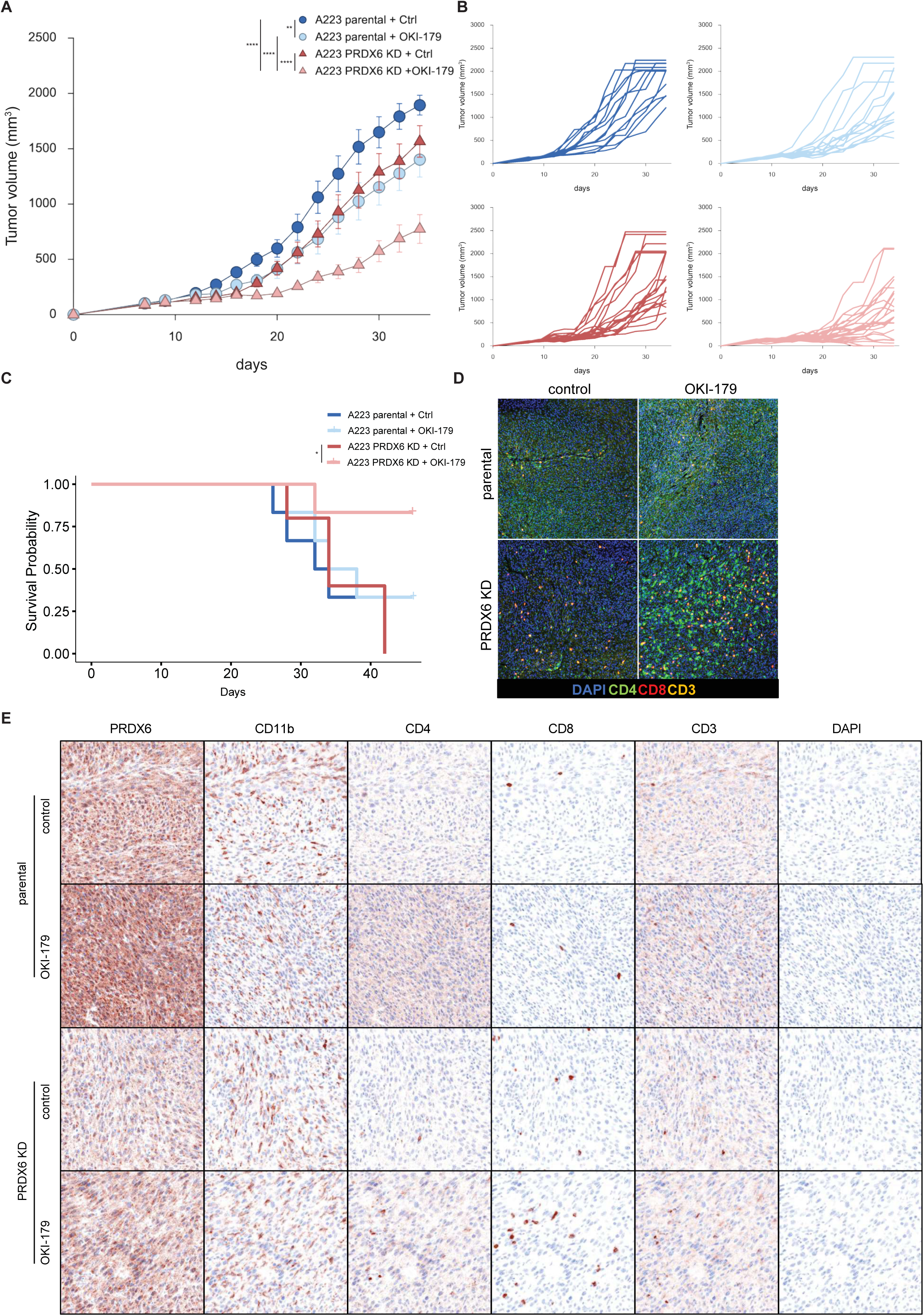
PRDX6 knockdown sensitizes largazole treatment *in vivo*. **(A)** PRDX6 KD enhances tumor growth inhibition by OKI-179 in A223 tumors. A223 parental or PRDX6 KD cells were implanted in the flank of WTB6 mice. After tumor volume reached ∼180 mm^3^ (A223 WT recipient mice were treated from day 12 onwards and PRDX6 KD recipient mice were treated from day 16 onwards) animals were treated with 60mg/kg OKI-179 or vehicle via oral gavage at 2 days interval for 9 doses. Tumor volume was measured and groups were compared on day 34. A223 parental + Ctrl, n=13; A223 parental + OKI-179, n=14; A223 PRDX6 KD + Ctrl, n=19; A223 PRDX6 KD + OKI-179, n=23. Different groups were compared using two-way ANOVA followed by Tukey’s multiple comparisons test. \*\**p*<0.01, \*\*\*\**p*<0.0001. **(B)** Individual tumor growth curves of recipients in different treatment groups. **(C)** Kaplan-Meier survival curve of recipient mice in different treatment groups. Groups were compared using log-rank tests. \**p*<0.05. **(D)** Representative immunohistochemical staining (DAPI, CD4, CD8 and CD3) of tumor tissues from A223 WT and PRDX6 KD mice with and without OKI-179 treatment. **(E)** Representative immunohistochemical staining (DAPI, PRDX6, CD11b, CD4, CD8 and CD3) of tumor tissues from A223 WT and PRDX6 KD mice with and without OKI-179 treatment.

### PRDX6 depletion enhances CD8 and CD4 T cell infiltration and reduces CD11b myeloid cells in tumors in response to OKI-179 treatment

Because our previous data suggest that PRDX6 KD cells treated with largazole exhibit a more ferroptosis-permissive, immunostimulatory state, the enhanced tumor growth inhibition by OKI-179 in A223 PRDX6 KD tumors could result from modulating anti-tumor immunity in addition to directly targeting tumor cells. To test this notion, we performed 7-color multiplex spectral imaging (MSI) with the Akoya system on tumor samples from experiments shown in Figure 6D and 6E. To profile the tumor and TME, we used a panel of antibodies including PRDX6, cytokeratin (CK5, tumor), CD3, CD8, CD4, CD11b plus DAPI. As expected, PRDX6 expression was significantly lower in PRDX6 KD tumor compared to WT (Figure 6E). In control WT A223, almost no T cells can be detected (Figure 6D and 6E). In PRDX6 KD A223 tumor, sporadic CD4 and CD8 cells can be seen (Figure 6D and 6E). OKI-179 treatment caused a much higher increase in CD4 and CD8 T cells and substantial decrease in CD11b positive myeloid cells in A223 PRDX6 KD tumors than in WT tumors (Figure 6E). This observation aligns closely with our cytokine array findings, which revealed significant reduction in macrophage-related cytokines (Figures 5E1, 5E2, and Supplemental Figures S9). These results suggest that reducing PRDX6 expression inhibits tumor growth and is associated with increased T-cell infiltration and modulation of the TME in response to OKI-179 treatment.

### Combined PRDX6 knockdown and OKI-179 enhances therapeutic efficacy of anti-PD-L1 immunotherapy

To further explore the therapeutic potential of combining PRDX6 inhibition and HDAC blockade with immune checkpoint inhibitors, we employed a syngeneic mouse model with A223 parental and PRDX6 KD tumors. Mice bearing either parental or PRDX6 KD tumors were treated with vehicle control, OKI-179, anti-PD-L1 antibody, or a combination of OKI-179 and anti-PD-L1 antibody (Figure 7A-7D). Survival analyses demonstrated that PRDX6 KD tumors exhibited significantly improved responses compared to WT tumors under identical treatments. Notably, the combined treatment of OKI-179 and anti-PD-L1 dramatically enhanced survival outcomes exclusively in mice with PRDX6 KD tumors, highlighting a strong combined treatment effect (Figure 7E and 7F).

**Figure 7:**
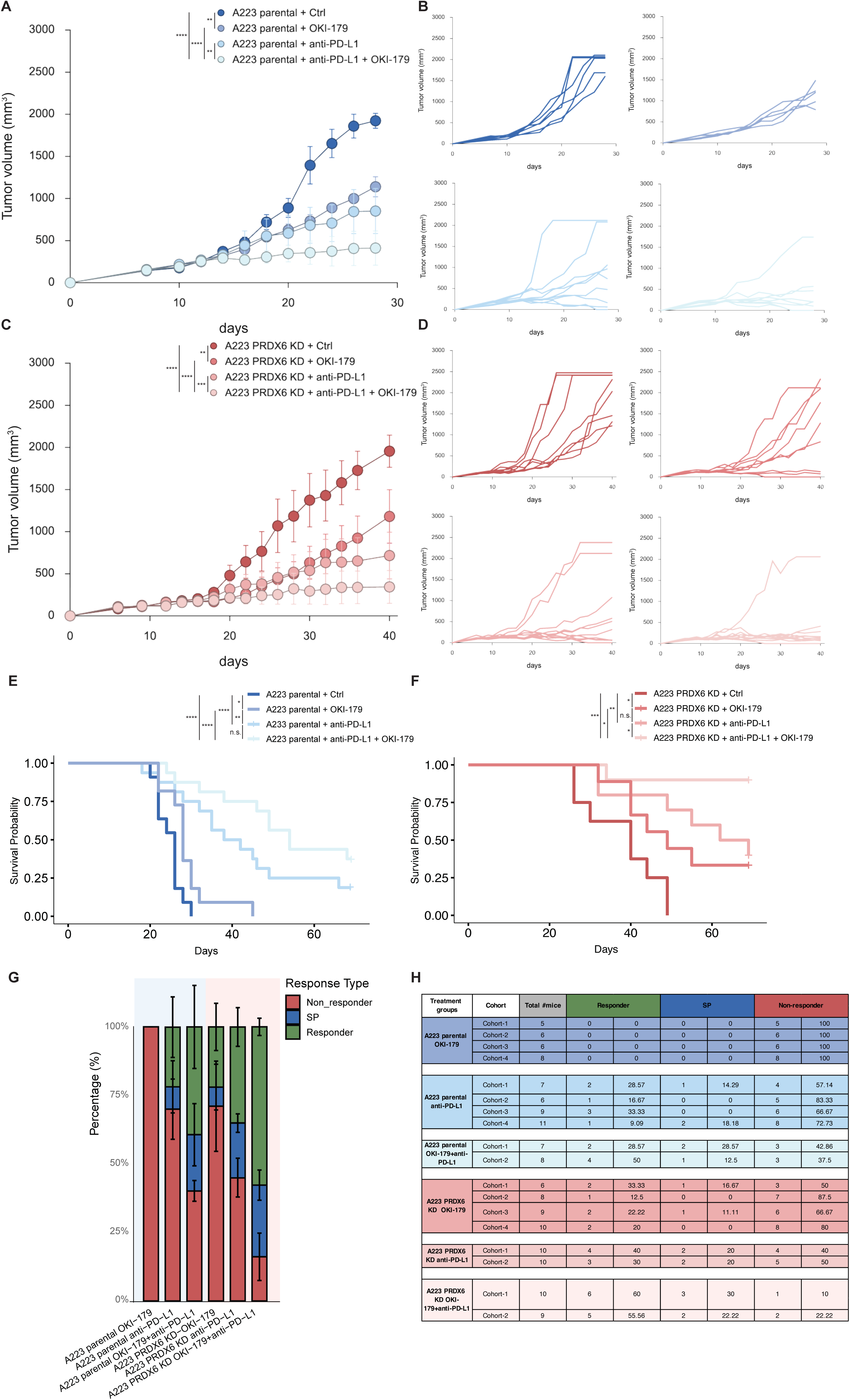
PRDX6 modulates immune checkpoint inhibitor response. Combination of anti-PD-L1/OKI-179 led to better outcomes in A223 PRDX6 KD cell line. **(A)** A223 parental cells were implanted in the flank of WTB6 mice and on day 12 tumor bearing mice were randomized into 4 groups, Control (n=6), OKI-179 (60 mg/kg/dose/2 days via oral gavage for 9 doses, n=5), anti-PD-L1 (200 µg/mouse/2days, i.p., for 3 doses, n=9), and OKI-179+anti-PD-L1 combination (n=8). Tumor volume was measured and groups were compared on day 28. **(C)** A223 PRDX6 KD cells were implanted in the flank of WTB6 mice and on day 16 tumor bearing mice were randomized into 4 groups, Control (n=8), OKI-179 (60 mg/kg/dose/2 days via oral gavage for 9 doses, n=9), anti-PD-L1 (200 µg/mouse/2days, i.p., for 3 doses, n=10), and OKI-179+anti-PD-L1 combination (n=10). Tumor volume was measured and groups were compared on day 40. Different groups were compared using two-way ANOVA followed by Tukey’s multiple comparisons test. \*\**p*<0.01, \*\*\**p*<0.001, \*\*\*\**p*<0.0001. **(B-D)** Individual tumor growth curves of recipients in different treatment groups of A223 parental and A223 PRDX6 KD cells. **(E, F)** Kaplan-Meier survival curves of recipient mice in different treatment groups A223 parental and A223 PRDX6 KD cells. Groups were compared using log-rank tests. \**p*<0.05, \*\**p*<0.01, \*\*\**p*<0.001, \*\*\*\**p*<0.0001, n.s. = not significant. **(G-H)** Combination of anti-PD-L1/OKI-179 led to a higher percentage of responders. A223 WT or PRDX6-KD cells were implanted at the flank of WT B6 mice and treated as described above. RCTV was calculated and compared for different treatment groups. Based on RCTV values, treatment recipients were divided into Responders (R, RCTV<0), Slow progressors (SP, 0<RCTV≤1.5), Non-responders (NR, RCTV>1.5). **(G)** Percentage of each response group (R, SP or NR) in different treatment groups. Different groups were compared using one-way ANOVA followed by Tukey’s multiple comparisons test. \**p*<0.05, \*\**p*<0.01, \*\*\**p*<0.001, \*\*\*\**p*<0.0001. **(H)** Summary of responses to different treatment regimens. For each treatment regimen, multiple cohorts of mice were used for independent experiments.

To quantitatively evaluate therapeutic responses, we calculated the relative change in tumor volume (RCTV) and classified mice as responders (R, RCTV<0), slow progressors (SP, 0<RCTV≤1.5), or non-responders (NR, RCTV>1.5). The combination treatment of anti-PD-L1 with OKI-179 in PRDX6 KD tumors resulted in the highest proportion of responders among all treatment groups, consistent with synergy or additive effects between PRDX6 deficiency and HDAC inhibition in potentiating checkpoint blockade (Figure 7G and 7H). Statistical analyses further supported significant differences in response profiles across treatment groups, underscoring the improved outcome of the combined regimen. Across multiple independent experiments, we observed reproducible enhancement of efficacy with combined PRDX6 inhibition and HDAC blockade, particularly in combination with anti-PD-L1. Taken together, these data indicate that PRDX6 deficiency downregulates GPX4 in squamous carcinoma, heightens ferroptosis induced by the HDAC inhibitor largazole, and reshapes the tumor microenvironment to promote T-cell infiltration and differentiation through multiple cytokine release. These findings support targeting PRDX6 alongside HDAC inhibition as a promising strategy to improve responses to immune checkpoint therapy (Supplemental figure S10).

## DISCUSSION

In this study, we identified Peroxiredoxin 6 (PRDX6) as a key mediator of resistance to the HDAC inhibitor largazole through a genome-wide CRISPR activation (CRISPRa) screen. We show that PRDX6 overexpression confers resistance to largazole-induced cell death independently of canonical HDAC target pathways. Conversely, PRDX6 depletion sensitizes cancer cells to largazole by elevating lipid peroxidation stress due to loss of PRDX6’s phospholipase A2 (PLA2) activity. This lipid peroxidation overload triggers ferroptosis and initiates immunogenic remodeling of the tumor microenvironment. Mechanistically, PRDX6 positively regulates GPX4, and its loss potentiates ferroptosis in part through GPX4 downregulation. TCGA analysis identified head and neck squamous cell carcinoma (HNSC) as particularly sensitive to alterations in PRDX6 expression, which we further validated in a syngeneic SCC model. In PRDX6-deficient SCC tumors, treatment with the clinical-stage HDAC inhibitor OKI-179 (largazole analog) significantly enhanced T-cell infiltration, upregulated pro-inflammatory cytokines and chemokines, and reduced immunosuppressive mediators such as G-CSF, GM-CSF, M-CSF, GDF-15, and osteopontin. Leveraging these immunogenic effects, we further demonstrated that combining OKI-179 with anti-PD-L1 immune checkpoint blockade markedly improved survival outcomes in mice bearing PRDX6-deficient tumors. Together, our findings suggest that targeting PRDX6 in combination with HDAC inhibition and immune checkpoint blockade may enhance antitumor immune responses and promote a more immunogenic tumor microenvironment.

Histone deacetylase (HDAC) inhibitors are a promising class of anticancer agents that exert their effects through epigenetic reprogramming, altering chromatin structure and transcriptional output to promote tumor suppression ^9,10,46^. By blocking HDAC catalytic activity, these compounds promote histone hyperacetylation, resulting in derepression of tumor suppressor genes and inhibition of oncogenic pathways. This leads to classical phenotypes such as cell-cycle arrest, apoptosis, and differentiation ^9,10,46,47^. While FDA-approved HDAC inhibitors including vorinostat, belinostat, panobinostat, and romidepsin, have shown efficacy in certain hematologic malignancies, their impact in solid tumors remains limited ^48^. The inhibition of proliferation and induction of apoptosis, while necessary, are insufficient to generate durable responses in most solid tumors. Recent studies suggested that tumor cell-intrinsic responses to HDACi, including modulating tumor immunogenicity and chemokines and cytokines that activate immune system, may have substantial roles in mediating the antitumor effects of HDACi ^11,49^. A critical deficiency in the field is a defined mechanism, linked to specific molecular targets or pathways, that converts HDACi-induced stress into sustained immunogenic pressure on tumors.

Through an unbiased CRISPRa screen identified PRDX6, and specifically its phospholipase A2 (PLA2) enzymatic function, as a key suppressor of largazole-induced cytotoxicity (Figures 1 and 2). While PRDX6 had not been previously implicated in HDAC inhibitor resistance, prior studies have linked PRDX6 deficiency ^50^ or HDAC inhibition ^51^ individually to ferroptosis. Our findings provide a mechanistic bridge between these observations by showing that largazole induces ferroptotic stress, and that PRDX6 loss intensifies this effect by disabling lipid hydroperoxide detoxification via its PLA2 activity (Figure 3). Our findings provide a mechanistic link between these observations: largazole triggers ferroptotic stress, and loss of PRDX6 amplifies this effect by impairing lipid hydroperoxide detoxification via its PLA2 function. Our results demonstrate that HDAC inhibitors such as largazole can engage ferroptosis management machinery when antioxidant defenses, such as PRDX6, are compromised.

A central regulator of ferroptosis is GPX4, a selenocysteine-containing enzyme that detoxifies lipid hydroperoxides ^52^. While most therapeutic strategies have focused on direct GPX4 inhibition (e.g., with RSL3), there is growing interest in targeting upstream regulators of GPX4 as alternative means to induce ferroptosis. PRDX6 has been proposed to act as a selenium carrier that facilitates GPX4 synthesis ^39–41^. Our data are consistent with this functional linkage: PRDX6 knockdown decreases GPX4 protein expression, and largazole further enhances this suppression. Moreover, our results indicate that this functional connection extends beyond selenium delivery and translational regulation. The coordinated expression of PRDX6 and GPX4 also occurs at the mRNA level, suggesting that PRDX6 modulation affects GPX4 through both transcriptional and post-transcriptional mechanisms. In light of this mechanistic insight, we examined patient samples from a phase II clinical trial of the class I selective HDAC inhibitor entinostat combined with nivolumab in unresectable or metastatic, previously treated pancreatic ductal adenocarcinoma^42^. The trial enrolled thirty patients, and RNA seq data were available for sixteen treated patients. Coordinated reductions in PRDX6 and GPX4 mRNA were observed after treatment, with the greatest decreases seen in the two patients with partial responses (Supplemental Figure S8B, S8C, S8D). Although the sample size is small and does not allow a definitive statistical conclusion, these findings raise the possibility that concurrent suppression of PRDX6 and GPX4 could serve as a biomarker for response to HDAC inhibitor and nivolumab therapy. Taken together, these results reveal a dual benefit of targeting PRDX6. Loss of PRDX6 sensitizes tumors to ferroptosis by impairing lipid detoxification, and at the same time it undermines GPX4 expression itself. While this mechanism appears to be conserved in human and murine cancer cells, additional studies are needed to determine how broadly it applies across tumor types and how best to exploit this vulnerability therapeutically.

One possible molecular mechanism that may explain the coordinated regulation of PRDX6 and GPX4 is that both PRDX6 and GPX4 have been shown to be regulated by SP1 transcription factor ^53–55^. We showed that largazole evicts SP1 from chromatin and suppresses SP1-dependent transcription ^56^. Class I HDAC inhibitors may activate ferroptosis response by targeting the antioxidant defense network through SP1 suppression. Thus, our findings uncover a previously unrecognized ferroptotic axis underlying largazole response and establish PRDX6 as a critical rheostat of HDACi-induced oxidative stress. Ferroptosis has recently attracted significant attention for its potential role in cancer immunotherapy, largely due to its immunogenic characteristics ^15,57,58^. Early-stage ferroptotic cells can release damage-associated molecular patterns (DAMPs), such as high-mobility group box 1 (HMGB1) and ATP, which stimulate dendritic cells and subsequently promote adaptive anti-tumor immune responses ^59–61^. However, the immunogenicity of ferroptosis appears to be context-dependent, influenced by factors including cancer type, the specific ferroptosis-inducing agents used, and interactions within the tumor microenvironment ^62,63^. Additionally, oxidized lipid species released during ferroptosis have been proposed to modulate its immunogenic potential, though their precise role remains incompletely understood ^64^. Consistent with the immunostimulatory nature of ferroptosis, we found that depletion of PRDX6 in SCC cells elevates lipid peroxidation and elicits pro-inflammatory and immunogenic gene expression signatures concurrent with a reduction in immunosuppressive factors upon largazole treatment. This shift in response profile may have contributed to reduction of CD11b+ myeloid cells and infiltration of CD8 and CD4 positive T cells to tumors due to enhanced tumor immunogenicity. Our analysis of the TCGA dataset reveals that overall patient survival is negatively correlated with PRDX6 expression in seven cancer types, with the most pronounced effect observed in HNSCC. Indeed, depletion of PRDX6 in mouse and human SCC lines significantly retards tumor cell proliferation *in vitro* and *in vivo* (Figure 4 and Figure 6). Head and neck cancers, particularly squamous cell carcinomas, are known to exhibit distinct reactive oxygen species (ROS) levels, with a subpopulation of ROS-Low cells showing enhanced cancer stemness and chemoresistance ^65,66^. It is tempting to speculate that PRDX6-dependent antioxidant systems may contribute to poor treatment response. Additionally, despite recent approval of pembrolizumab, an anti-programmed cell death protein 1 (PD-1) antibody by the FDA as a first-line treatment for recurrent/metastatic HNSCC, the majority of patients do not respond to this treatment, and overall survival remains limited with few long-term survivors ^67^. Our findings reveal that combining PRDX6 deficiency and HDAC inhibition with anti-PD-L1 immune checkpoint blockade markedly improves therapeutic responses in a head neck cancer model (Figure 7) suggests that high levels of PRDX6 also reduce tumor immunogenicity and contribute to poor response rate. Given the highly context-dependent nature of ferroptosis-driven immunogenicity, it is crucial to test this notion in different HNSCC derived tumor models and patient derived tumor models.

Our results suggest that pairing HDAC inhibitors with PRDX6 inhibitors, or using both with immune checkpoint inhibitors, may produce more durable antitumor responses and convert epigenetic stress into immunogenic, irreversible tumor elimination. The natural product Withangulatin A (WA) inhibits PRDX6 through irreversible covalent modification of the cysteine 47 residue and suppresses proliferation of non-small cell lung cancer cells ^68^. However, WA also reacts with other protein thiols, including thioredoxin reductase, PRDX1 and Nrf2, complicating attribution of its antitumor mechanism ^69^. MJ33, an analog of a phospholipid substrate intermediate, blocks the iPLA2 activity of PRDX6 *in vitro* (IC50 ∼300 nM) and suppresses LPS induced PRDX6 dependent acute lung injury when formulated in unilamellar liposomes ^32,70^. Limitations in bioavailability and biodistribution of the lipid analog MJ33 make it impractical for combination studies with OKI-179 or anti PD L1 at present. Future studies should prioritize discovery of more potent, selective and drug-like PRDX6 small molecule inhibitors suitable for systemic delivery, enabling rigorous combination testing and definitive validation of the therapeutic benefit of targeting PRDX6 in cancer. Based on the mechanism we uncovered, such inhibitors are expected to impair PRDX6-mediated lipid detoxification and GPX4 activity, providing a clear rationale for combination strategies.

In summary, our findings illuminate a previously unrecognized regulatory relationship between PRDX6 and GPX4 in ferroptosis resistance. By linking clinical outcomes, CRISPRa dependency, gene expression, and functional assays, we provide evidence that PRDX6 modulates ferroptosis susceptibility in squamous cancer models and may represent a vulnerability when combined with HDAC inhibition. Our work points to a broader conceptual advance: therapeutic strategies that combine HDAC inhibition with ferroptosis induction have the potential to reshape the tumor microenvironment, unlock antitumor immunity, and overcome the inherent limitations of cytostatic therapies.

## ACKNOWLEDGMENTS

We thank members of Wang and Liu laboratories for discussion and suggestions and Dr. Yadira Soto-Feliciano for critical reading of the manuscript. We also want to thank Anthony Piscopio and Nicolas Saccomano of OnKure Therapeutics for OKI-179. This work was supported by grants from National Institutes of Health R01GM144749 (Liu) and R01CA229174A1 (Wang). This work was also supported by a Developmental Research Award from CO HNC SPORE grant (P50CA261605) (Liu and Wang), an ABNexus award (Liu and Pitts) and a Milheim Foundation Pilot Grant (Liu). N.J. is supported by a predoctoral training grant from NIGMS (T32GM08759). The ImageXpress MicroXL was supported by a NCRR grant S10RR026680, and FACSAria was supported by S10OD021601, and Opera Phenix imaging system by S10OD025072 from NIH.

## AUTHOR CONTRIBUTIONS

Z.L., J.J., N.J., G.S., J.W., and X.L. designed the study and analyzed the data. Z.L., J.J., N.J., P.S., G.S., D.S., and X.L. performed experimental studies. Z.L., J.J., N.J., J. W., and X.L. wrote the manuscript in consultation with inputs from all authors.

## DECLARATION OF INTERESTS

X.L. is a co-founder of OnKure Therapeutics, Inc and Vesicle Therapeutics, Inc and owns equities in both companies. Neither of these companies were involved in the experimental design or funding this study.

## FIGURE LEGEND

**Supplemental Figure 1.** HDAC inhibition testing for hits from the CRISPRa screen. **(A)** Validation of ASS1 overexpression, ASS1 knockdown and PRDX6 overexpression in HCT116 cells. GAPDH was used as a loading control. **(B)** Validation of screening results using cells with individual gene overexpression or knockdown. HCT116 WT cells with indicated gene overexpression or knockdown using cDNA or shRNA were treated with 100 nM largazole over 12 days. After treatment, cells were fixed with 4% formaldehyde in PBS and stained with 0.05% crystal violet. A three day DMSO treatment of HCT116 WT cells was used as a natural growth control. Images were inverted into grayscale format to maximize the visibility. **(C)** Quantification of Figure S1B. Crystal violet stained samples were solubilized with the solubilization buffer and absorbance was measured at 570 nm. Data are mean ± SD. Welch’s one-way ANOVA with Games-Howell multiple comparisons test was used for statistical analysis. \*\*\*\**p* < 0.0001. **(D)** Western blot for pan-H4ac, H3K9ac and H3K27ac expression levels in HCT116 WT, ASS1 and PRDX6 OE cell lines. Cells were treated with largazole at the indicated concentrations for 2 hours and proceeded for Western blotting with the indicated makers. GAPDH was used as a loading control.

**Supplemental Figure 2.** PRDX6 knockdown sensitizes HCT116 cells to ferroptosis. **(A-B)** Dose-response matrix heatmaps for HCT116 WT and PRDX6 KD cells with largazole + erastin2. **(C-D)** Dose response matrix heatmaps for HCT116 WT and PRDX6 KD treated with largazole + RSL3. **(E-F)** 3d heatmaps of synergy HSA score for HCT116 WT and PRDX6 KD cells with largazole + RSL3.

**Supplemental Figure 3.** TCGA analysis reveals PRDX6 is a critical maker for survival probability across cancers within the PRDX family. **(A)** Survival analysis of the PRDX family in 33 TCGA tumor types. Heatmap displays the log2 hazard ratios (HR). Statistically significant associations (p < 0.05) are indicated with an asterisk (*). **(B)** 5-year Kaplan-Meier curves for overall survival from TCGA data for PRDX6. Patients were stratified into High-PRDX (red) and Low-PRDX (blue) groups based on the median cutoff. The shaded area represents the 95% confidence intervals. The log-rank test *p*-value < 0.05 for all curves. **HNSC** = head and neck squamous carcinoma. **ACC** = Adrenocortical carcinoma. **BRCA** = Breast invasive carcinoma. **KIRP** = Kidney renal papillary carcinoma. **LGG** = Brain lower grade glioma. **LUAD** = Lung adenocarcinoma. **UCEC** = Uterine corpus endometrial carcinoma.

**Supplemental Figure 4.** TCGA analysis reveals PRDX6 is a critical maker for survival probability of specific cancers among the PRDX family. 5-year Kaplan-Meier curves for overall survival from TCGA data for PRDX1 **(A)** and PRDX5 **(B).** Patients stratified into High-PRDX (red) and Low-PRDX (blue) groups based on median cutoff. The shaded area represents the 95% confidence interval. The log-rank test p-value < 0.05 for all curves. **HNSC** = head and neck squamous carcinoma. **ACC** = Adrenocortical carcinoma. **BRCA** = Breast invasive carcinoma. **KIRP** = Kidney renal papillary carcinoma. **LGG** = Brain lower grade glioma. **LUAD** = Lung adenocarcinoma. **UCEC** = Uterine corpus endometrial carcinoma.

**Supplemental Figure 5.** PRDX6 KD sensitizes the head and neck squamous carcinoma cell line FaDu to ferroptosis with largazole treatment. (**A** and **C**) After the indicated treatments, both FaDu shNT and PRDX6 KD cells were stained with 1 µM FerroOrange (Dojindo F374-10) or 1 µM LiperFluo (Dojindo L248-10) in complete FluoroBrite DMEM (10% FBS, 1% GlutaMAX, 1% pen/strep) and costained with 10 µg/mL Hoechst 33342 for 30 minutes. Cells were seeded in Revvity Phenoplate Ultra 96-well plates at 8,000–10,000 cells/well to ensure consistent confluency, rinsed with D-PBS, and imaged on the Revvity Opera Phenix High-Content Screening System using the 20X water objective. Imaging channels used were 561–584/25 nm for FerroOrange and 488–525/50 nm for LiperFluo, with standardized exposure, gain, and laser settings. (**B** and **C**) Hoechst-based nuclear segmentation and algorithmically defined perinuclear masks enabled single-cell quantification of cytoplasmic fluorescence in Harmony 5.2 software. Cells with high Fe²⁺ or lipid peroxidation were identified using Otsu’s threshold from DMSO-treated shNT controls and reported as the percentage of high-signal cells per well. Bars represent mean ± SEM from biological replicates. Statistical significance was determined by two-way ANOVA with Tukey’s post hoc test. \**p* < 0.05, \*\**p* < 0.01, n.s. = not significant. (**E**–**F**) Heatmap of the dose–response matrix for FaDu shNT and PRDX6 KD cells with largazole + RSL3. (**G**–**H**) Heatmap of the synergy HSA score for FaDu shNT and PRDX6 KD cells with largazole + RSL3.

**Supplemental Figure 6.** PRDX6 KD sensitizes mouse squamous carcinoma cells to ferroptosis. (**A** and **C**) After the indicated treatments, both A223 shNT and PRDX6 KD cells were stained with 1 µM FerroOrange (Dojindo F374-10) or 1 µM LiperFluo (Dojindo L248-10) in complete FluoroBrite DMEM (10% FBS, 1% GlutaMAX, 1% pen/strep) and costained with 10 µg/mL Hoechst 33342 for 30 minutes. Cells were seeded in Revvity Phenoplate Ultra 96-well plates at 8,000–10,000 cells/well to ensure consistent confluency, rinsed with D-PBS, and imaged on the Revvity Opera Phenix High-Content Screening System using the 20X water objective. Imaging channels used were 561–584/25 nm for FerroOrange and 488–525/50 nm for LiperFluo, with standardized exposure, gain, and laser settings. (**B** and **C**) Hoechst-based nuclear segmentation and algorithmically defined perinuclear masks enabled single-cell quantification of cytoplasmic fluorescence in Harmony 5.2 software. Cells with high Fe²⁺ or lipid peroxidation were identified using Otsu’s threshold from DMSO-treated shNT controls and reported as the percentage of high-signal cells per well. Bars represent mean ± SEM from biological replicates. Statistical significance was determined by two-way ANOVA with Tukey’s post hoc test. \**p* < 0.05, \*\*\**p* < 0.001, \*\*\*\**p* < 0.0001, n.s. = not significant. (**E**–**F**) Heatmap of the dose–response matrix for A223 shNT and PRDX6 KD cells with largazole + erastin2. (**G**–**H**) Heatmap of the synergy HSA score for A223 shNT and PRDX6 KD cells with largazole + erastin2.

**Supplemental Figure 7.** Cross-correlation of PRDX6 and GPX4 in HNSC based on analysis across multiple databases. **(A-D)** Perturbation effects of PRDX6 in different cell lines. Data were provided and visualized by DepMap using the CRISPR dataset (DepMap Public 25Q2+score, Chronos). **(E-H)** Co-expression analysis of PRDX6 and GPX4 in HNSC. Multiple datasets were used (S7E: TCGA, Nature 2015; S7F: TCGA, Firehorse Legacy; S7G: TCGA, GDC; S7H: TCGA PanCancer Atlas). Datasets were analyzed and visualized using cBioportal.

**Supplemental Figure 8.** HDAC inhibitor co-regulates PRDX6 and GPX4 in patient samples from a phase II clincal trial ^42^. (**A**) Overall expression levels of GPX4 and PRDX6 before and after entinostat treatment for each patient from published data (GSE248014). Downregulation of either gene post-treatment is indicated by a blue background, while upregulation is indicated by a red background. For each patient, concurrent upregulation of both genes is labeled in red, concurrent downregulation in green, and divergent regulation in black. (**B**) GPX4 and PRDX6 expression changes upon entinostat treatment across individual patients. Slope plots illustrate GPX4 and PRDX6 levels at baseline and following treatment. Each line connects paired measurements from the same patient. Green circles indicate a decrease in expression post-treatment, while red circles indicate an increase. (**C–D**) Correlation between GPX4 and PRDX family members before treatment (**C**) and after one cycle of entinostat (C1D1) (**D**). Scatter plots depict relationships between GPX4 expression and that of PRDX2, PRDX4, and PRDX6 across 16 patient samples. Each point represents a patient. Pearson correlation coefficients (r) and p-values are shown. Linear regression lines with 95% confidence intervals are included. Patient 18 and 39 are highlighted in red.

**Supplemental Figure 9.** Additional cytokine results for A223 parental and PRDX6 KD cells with largazole treatment. **(A)** Cytokine array shows elevated inflammatory response induced by largazole treatment in A223 PRDX6 KD cells. A223 parental and PRDX6 KD cells were seeded into a 6-well plate with 2×10^6^ cells in each well. After 24 hours incubation, DMEM was replaced by 1 mL of fresh DMEM with DMSO or 100 nM largazole for 18 hours. After treatment, each supernatant was collected and expression level of each cytokine was measured with Proteome Profiler™ Array Mouse XL Cytokine Array. **(B)** Quantification of Panel A. Intensity of each cytokine spot was measured and a Two-way ANOVA with Tukey multiple comparisons test was used for statistical analysis. \*\**p* < 0.01, \*\*\**p* < 0.001, n.s. = not significant.

**Supplemental Figure 10.** Schematic model of PRDX6-mediated regulation of ferroptosis and antitumor immunity. PRDX6 acts as a redox gatekeeper that suppresses lipid peroxidation and maintains redox homeostasis, thereby protecting tumor cells from ferroptotic cell death induced by the HDAC inhibitor largazole. Loss of PRDX6 downregulates GPX4 expression, sensitizing squamous carcinoma cells to ferroptosis and creating a metabolic vulnerability that synergizes with HDAC inhibition. This ferroptosis-sensitized state alters the cytokine landscape of the tumor microenvironment, promoting enhanced T-cell infiltration, differentiation, and effector function.

## CONTACT FOR REAGENT AND RESOURCE SHARING

Additional information and requests for resources and reagents should be directed to and will be fulfilled by the Lead Contact, Dr. Xuedong Liu.

## EXPERIMENTAL MODEL AND SUBJECT DETAILS

### Key resources table

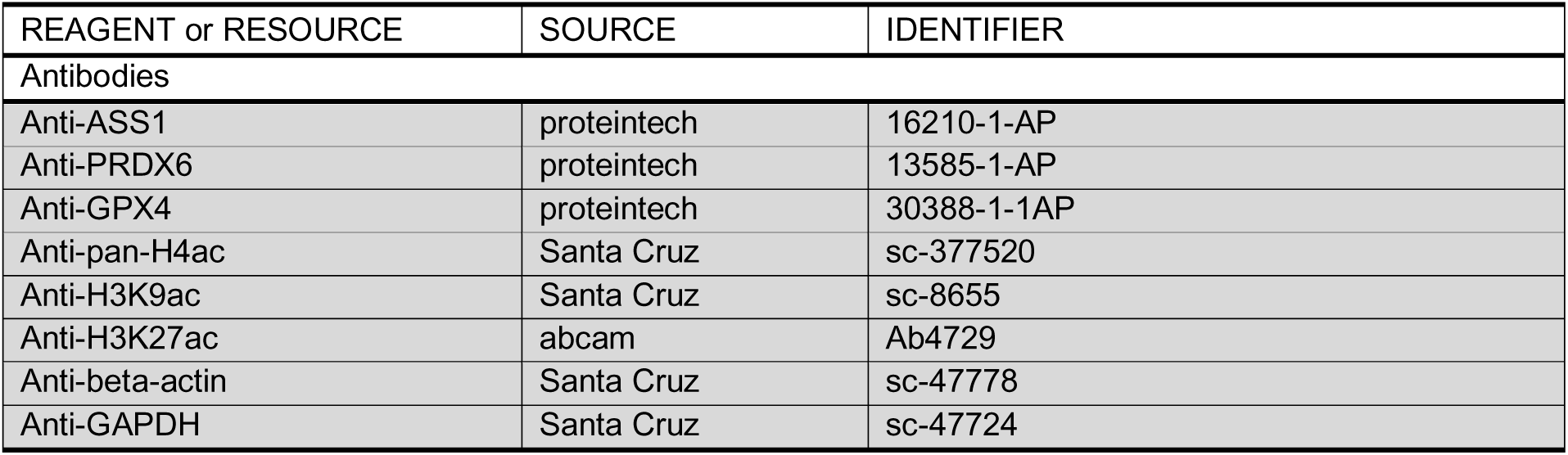
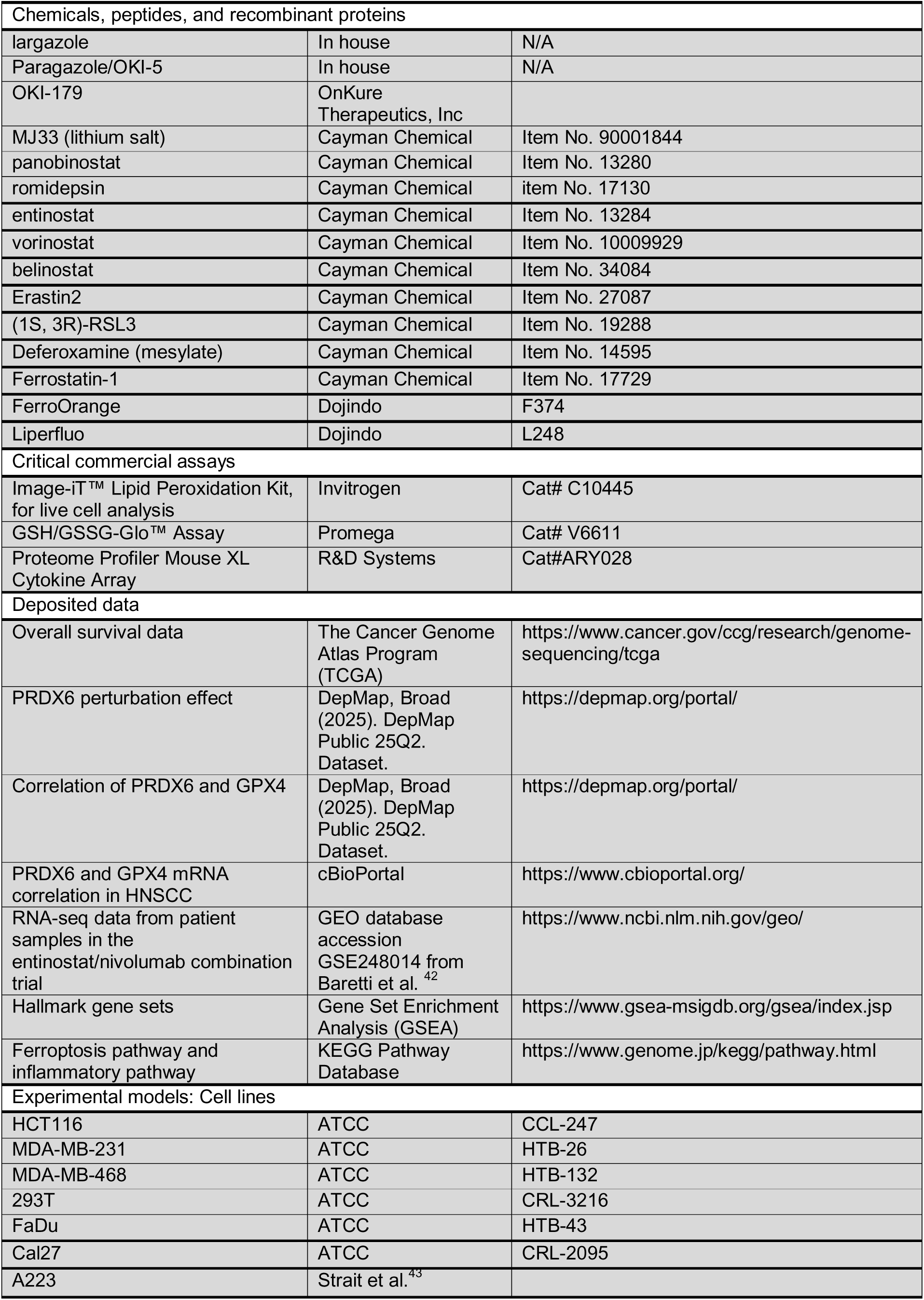
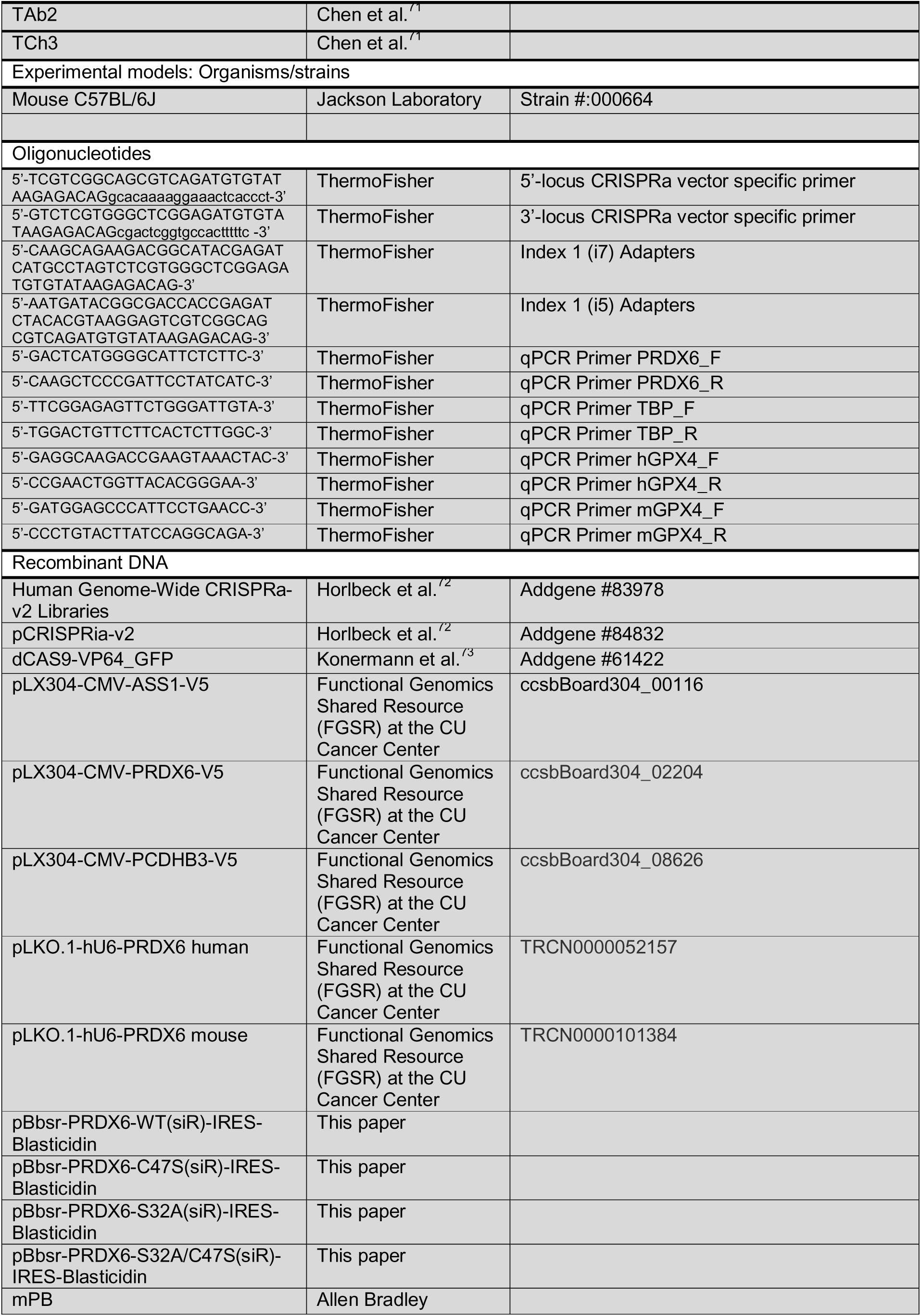
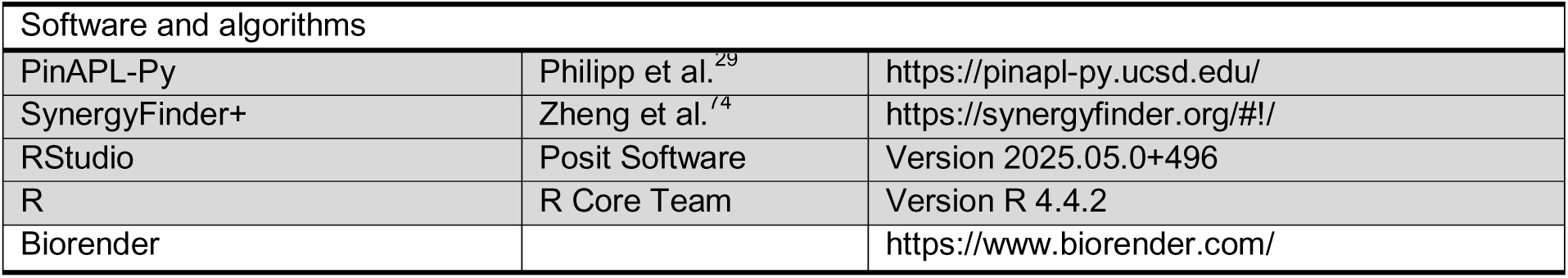

### Cell lines and cell culture

The human colorectal carcinoma (HCT116), triple-negative breast adenocarcinoma (MDA-MB-231 and MDA-MB-468) and the embryonic kidney epithelial (293T) cell lines were acquired from ATCC. The murine A223 was derived from a Smad4 deficiency squamous tumor and has been shown to be a suitable animal model for HNSCC ^43–45^. TAb2 and TCh3 cell lines were generated from K15.CrePR1(+)p53^f/f^PIK3CA^c/c^ mouse as described previously ^43,71^. All cells were grown in Dubelcco’s modified Eagle medium (DMEM) supplemented with 10% fetal bovine serum (FBS), 1% penicillin-streptomycin, and 1% GlutaMAX (Invitrogen). Cell lines were maintained at 37°C with 5% CO_2_ and tested routinely for mycoplasma contamination.

## METHOD DETAILS

### 100x Library Coverage Genome-scale CRISPRa Screen and Data Analysis

The genome-scale screen was conducted similar to the protocol described by Weissman Lab (https://weissman.wi.mit.edu/crispr/). The CRISPRa v2 library (Addgene, #83978) were transduced into HCT116 CRISPRa cells, at MOI < 0.5 (percentage of transduced cells 2 days after transduction: 40-50%, checked by BFP fluorescence ratio using BD FACSCelesta). Two days after transduction, cells were selected with 2 μg/mL puromycin for 2 days to reach the point where transduced cells accounted for 95-99% of the population. After the selection, 100x library coverage (25×10^6^-30×10^6^ cells) were harvested and the remaining cells were split into two populations for DMSO-treated and largazole-treated growth. Two populations were cultured in 15 cm culture dishes with seeding density 5×10^6^ for each dish at the beginning. DMSO-treated and largazole-treated populations were harvested after each dish reached cell density at 20×10^6^ (largazole-treated cell population was treated with 100 nM largazole, and additional pulses of largazole treatment were performed every 3 days). Genomic DNA was isolated from frozen cell pellets and the sgRNA-encoding region was enriched, amplified, and processed for sequencing on an Illumina NovaSeq6000.

Sequenced libraries were analyzed using PinAPL-Py (www.pinapl-py.ucsd.edu) ^29^. The project parameters were defined in the following way: the screen type was set as “enrichment”, the 5’-adapter sequence was defined as ‘CCCTTGGAGAACCACCTTGTTN’, and the reference library was obtained from the Weissman Lab repository at Addgene (https://www.addgene.org/pooled-library/weissman-human-crispra-v2/). We used the sgRNA ranking files and removed all data points where the treated and control output contained zero counts and used R version 4.1.0 to generate the plots.

### Focused CRISPRa Library Generation and Screen

Oligonucleotide pools were retrieved from 100x genome-scale screening sequence samples and cloned into the sgRNA expression vector pCRISPRia-v2 (Addgene, #84832). Focused library screen and analysis were conducted in the same manner as described previously in 100x Library Coverage Genome-scale CRISPRa Screen and Data Analysis

### DNA Transfection and Viral Infection

Lentivirus was produced by transfecting HEK293T cells with standard packaging vectors using 1mg/mL PEI (Polysciences, Inc., 1:5 μg DNA: μg PEI). Cultural medium was replaced 8 hours after transfection, and supernatant was harvested 2 days after medium replacement. Lentivirus supernatant was further filtered by 0.2 μm PVDF filters (Thermo Scientific™ Nalgene™) and mixed with polybrene (Sigma-Aldrich®, 10 μg/mL) before infection.

### shRNA-Mediated Knockdown and cDNA-Mediated Overexpression

To generate ASS1 KD and PRDX6 KD cell lines, WT HCT116, MDA-MB-231, MDA-MB-468, A223, TAb2 and TCh3 cells were transduced with lentiviral pLOK.1 vector expressing shRNA from an U6 promoter and selected for puromycin-positive cells using puromycin (TOCRIS®, 2 μg/mL).

To generate HCT116 cell lines stably expressing ASS1, PCED1B, R3HDM2, ENO2 and PRDX6, WT HCT116 cells were transduced with lentiviral PXL304 vector expressing indicated gene from an CMV promoter with a C-terminal V5 tag (ccsbBroad304, #00116, #04547, #15744, #08626, #06161, #02204) and selected for blasticidin-positive cells using blasticidin (Gibco™, 10 μg/mL). For LRRK, Human cDNA expression clone for LRRK1 was constructed by subcloning human LRRK1 cDNA from MGC premier ORF clone for LRRK1 (BC172446, Transomic) via Gateway cloning into pBbsr expression vector.

### Cell Viability Assay

3×10^5^ cells were seeded into each well of 6-well plate. After 24 hours of incubation, DMEM was replaced by fresh DMEM with 100 nM largazole. Additional pulses of largazole treatment were performed every 3 days. After total 12 days treatment, cells were washed with PBS twice and incubated with 1 mL of 4% formaldehyde in PBS at room temperature, 100 rpm shaking for 10 minutes (Thermo, MaxQ 2000). Cells were washed again with PBS after incubation and incubated with 1 mL of 0.05% crystal violet staining buffer (250 mL methanol, 750 mL MiliQ water, 0.5g crystal violet) at room temperature, with 100 rpm shaking for 10 minutes. Cells were washed with DI water and air dried for 24 hours. Dried stained samples were captured by a camera (Nikon D5600) and resolubilized using 6 mL of solubilization buffer (400 mL ethanol, 100 mL methanol, 100 mL MiliQ water) under 4°C, with 100 rpm shaking for 1 hour. Resolubilized samples were further quantified by plate reader (absorbance 530 nm, Molecular Devices SpectraMAX iD3).

### Western Blot

Cells with different treatments were harvested and lysed in the RIPA buffer. Cell lysates were separated on SDS-PAGE gels and transferred to nitrocellulose membranes (Amersham™ Protran™, 0.2 μm). After blocking nonspecific bindings with blocking buffer (5% non-fat milk in TBST) for 45 minutes at room temperature, membranes were blotted with primary antibodies ASS1(proteintech, 16210-1-AP), PRDX6 (proteintech, 13585-1-AP), H4ac (Santa Cruz, sc-377520), H3K9ac (Santz Cruz, sc-8655), H3K27ac (abcam, ab4729) and GAPDH (Santa Cruz, sc-47724) 1.5 hours at room temperature. After, membranes were incubated with HRP-conjugated donkey anti-rabbit/sheep anti-mouse secondary antibody (Cytiva, NA9311ML) for 1 hour at room temperature. The protein bands were developed with SuperSingal™ west dura extended duration substrate (Thermo, lot#X8340749) and detected by ImageQuant™ LAS 4000 (GE healthcare)

### PRDX6 Vectors Construction and PRDX6 Rescue Cell Lines Generation

Wild type and mutant human PRDX6 coding sequences were codon optimized and synthesized by Twist Biosciences. They were digested with BglII and NotI and subcloned into pBbsr-DEST vector digested with BamHI and NotI and sequencing verified. The codon optimized coding sequences are designed to have mismatches to PRDX6 siRNA used in this study and immune to RNAi. To generate HCT116 PRDX6 KD with PRDX6 WT, PRDX6 S32A, PRDX6 C47S and PRDX6 S32A C47S rescue cell lines, HCT116 PRDX6 KD cells were transduced with PRDX6-WT, PRDX6-S32A, PRDX6-C47S, PRDX6-S32A-C47S along with a piggy-back expression vector mPB and selected for blasticidin resistance clones.

### Largazole and MJ33 Combination Assay

HCT116 WT cells (3×10^5^) were seeded into each well in 6-well plates. After 24 hours of incubation, DMEM was replaced by fresh DMEM with 100 nM largazole. Additional pulses of largazole treatment were performed every 3 days. Cells were washed twice with PBS after total 9 days treatment, and each well was treated either with 2 mL of 100 nM largazole, 10 μM MJ33 or 100 nM largazole + 10 μM MJ33 for 3 days. Cells were collected and quantified in cell viability assay after 3 days treatment.

### Cell Viability Assays for Drug IC_50_

Each type of cells was seeded into 4 lanes x 12 wells of 96-well plate with 4000 cells seeded into each well. 12 hours after seeding, DMEM was replaced by fresh DMEM with indicated drug in serial-diluted manner, 12 different concentrations and 4 repeats, and cells were treated for 3 days. After treatment, for each well, cells were washed by 100 μL D-PBS twice and incubated with 50 μL of 4% formaldehyde in D-PBS under room temperature, 100 rpm shaking for 10 minutes (Thermo, MaxQ 2000). After 10 minutes shaking, cells were washed again with 100 μL D-PBS and incubated with 50 μL of 0.05% crystal violet staining buffer (250 mL methanol, 750 mL MiliQ water, 0.5g crystal violet) under room temperature, 100 rpm shaking for 10 minutes. After shaking, cells were washed with DI water and air dried for 24 hours. Dried stained samples were resolubilized using 150 μL solubilization buffer (400 mL ethanol, 100 mL methanol, 100 mL MiliQ water) under 4°C, 100 rpm shaking for 1 hour. Resolubilized samples were further quantified by plate reader to measure the intensity of solubilized crystal staining (absorbance 530 nm, Molecular Devices SpectraMAX iD3).

### GSH/GSSG-Glo™ Assay Normalized with CellTiter-Fluor™ Cell Viability Assay

Experiments were performed based on the protocol provided by Promega. Briefly, each cell line (4000 cells) was seeded into each well in 96-well plates. After 24 hours of incubation, DMEM was replaced by fresh DMEM with indicated concentrations of drugs and incubated for indicated duration. After the treatment, 20 µL of CellTiter-Fluor™ Reagent (prepared as 10 µL substrate in 2 mL Assay Buffer) was added to all wells, and mixed briefly by orbital shaking. Plates were incubated for at least 30 minutes at 37°C. After incubation, resulting fluorescence was measured using a fluorometer (380–400 nm_Ex_/505 nm_Em_, Molecular Devices SpectraMAX iD3). After the measurement, liquid from each well was removed and 50 µL per well of Total Glutathione Lysis Reagent or Oxidized Glutathione Lysis Reagent was added into wells. Plates were incubated and mixed briefly by orbital shaking for 5 minutes. After incubation, 50 µL of Luciferin Generation Reagent was added to each well and plates were incubated and mixed briefly by orbital shaking for 30 minutes. After incubation, 100 µL of Luciferin Detection Reagent was added to each well and plated were gently mixed by shaking and incubated at room temperature for 15 minutes in the dark. Luminescence was measured by the plate reader (Molecular Devices SpectraMAX iD3). For analysis, luminescence readings from GSH/GSSG-Glo™ Assay were normalized by fluorescence readings from CellTiter-Fluor™ Cell Viability Assay for each well. After normalization, luminescence readings were converted into concentrations based on the standard curve provided in the kit.

### FerroOrange Staining and Imaging for Iron Accumulation

Live cells were stained according to the manufacture’s parameters in the <u>FerroOrange F374</u> <u>Product Manual</u> (Dojindo F374-10). Stock solution was 24ug of dry product resuspended in 35uL DMSO to create a 1mM stock. Stock was further diluted to a 1uM working solution in full Fluorobrite DMEM (Gibco), complete with 10% FBS, 1% glutamax, and 1% pen/strep to maintain consistency to other culture conditions and experiments. Cells were seeded in Revvity/Perkin-Elmer 96-well Phenoplate Ultra (Revvity 6055302) at a density of 8000-10000 cells/well per cell line growth rates to achieve equivalent confluency at the time of drug application and imaging. Cells were rinsed with D-PBS once and dyed 30min prior to imaging costained with 10 µg/mL Hoechst 33342 (Thermo), dye media was not removed prior to imaging per manufacturer recommendations. Measurements were acquired with Revvity Opera Phenix Confocal High Content Screening utilizing the 20X Water Objective, channels HOECHST 33342 and 561-584/25 set to standard exposure, gain and laser power parameters.

### Lipids Peroxidation Detection Assay and Analysis

Each cell line (4000 cells) was seeded into each well in 96-well plates. After 24 hours of incubation, DMEM was replaced by fresh DMEM with indicated concentrations of drugs and incubated for indicated duration. After the treatment, cells were washed with PBS three times, then fixed with 4% paraformaldehyde in PBS by incubating in the dark under constant gentle shaking for 15 minutes. After fixation, fixed cells were gently washed with PBS three times, and stained by 100 µL of PBS with 10 µM Image-iT® Lipid Peroxidation Sensor (ThermoFisher) and 10 µg/mL Hoechst 33342 (Thermo) in each well for 3 days in the dark under 4°C. After staining, cells were washed three times with PBS and imaged with FITC, TRITC and DAPI channels using a microscope (PerkinElmer, Opera Phenix™). For imaging analysis, Harmony® was used for automatic quantification. In general, cells in each image were recognized by their nuclear maker, and fluorescent intensity of FITC and TRITC were quantified for each cell. The FITC/TRITC ratio of fifty cells from each image were randomly selected for analysis and presentation purposes.

### Liperfluo Staining and Imaging for Lipid Peroxidation

Live cells were stained according to the manufacture’s parameters in the Liperfluo L248 Product Manual (Dojindo L248-10). Stock solution was 50ug of dry product resuspended in 60uL DMSO to create a 1mM stock. Stock was further diluted to a 1uM working solution in full Fluorobrite DMEM (Gibco), complete with 10% FBS, 1% glutamax, and 1% pen/strep to maintain consistency to other culture conditions and experiments. Cells were seeded in Revvity/Perkin-Elmer 96-well Phenoplate Ultra (Revvity 6055302) at a density of 8000-10000 cells/well per cell line growth rates to achieve equivalent confluency at the time of drug application and imaging. Cells were rinsed with D-PBS once and dyed 30min prior to imaging costained with 10 µg/mL Hoechst 33342 (Thermo), dye media was removed prior to imaging per manufacturer recommendations. Measurements were acquired with Revvity Opera Phenix Confocal High Content Screening utilizing the 20X Water Objective, channels HOECHST 33342 and 488-525/50 set to standard exposure, gain and laser power parameters.

### Single Cell Analysis of FerroOrange and LiperFluo Measurements

Excess intracellular iron and lipid peroxidation were quantified using FerroOrange and LiperFluo (Dojindo) on a Revvity Opera Phenix High-Content Imaging System with Harmony 5.2 software. A custom analysis pipeline was developed in collaboration with a Revvity image analyst. Nuclei were segmented using Hoechst 33342 (Thermo Fisher Scientific) fluorescence, and a proportional perinuclear mask was algorithmically generated for each cell to capture cytoplasmic signal. Border cells at the edges of each field were excluded to prevent duplicate counting across image frames. Within each perinuclear mask, the total integrated fluorescence intensity of FerroOrange or LiperFluo was calculated. This measure provided both global and punctate signal quantification on a per-cell basis. “High” iron or lipid peroxidation thresholds were defined using Otsu’s method from control wells (shNT cells treated with DMSO), and the percentage of high-signal cells per well was determined as:

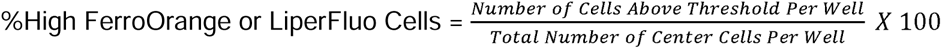

This approach enabled unbiased, cell-by-cell quantification of excess intracellular Fe²⁺ and lipid peroxidation levels.

### Synergism Analysis with SynergyFinder+

For each cell line, shNT and PRDX6 KD cells were plated in 961zwell plates in an 8 × 8 matrix layout (8 concentrations of largazole × 8 concentrations of erastin2/RSL3), at 8,000 cells per well. After ∼12 hours to allow cell adhesion, the medium was replaced with DMEM containing serial dilutions of largazole and erastin2/RSL3 in all pairwise combinations, and cells were incubated for 72 hours. Following treatment, wells were washed twice with PBS, fixed in 4% formaldehyde, stained with 0.05% crystal violet, and dye was solubilized; absorbance was then read at 530 nm. Untreated wells (no drug) were used as normalization controls to compute percent viability. Synergy computation and visualization were performed using SynergyFinder+ and HSA model was used for visualization and interpretations. Positive scores indicate synergistic interaction, negative scores indicate antagonism, and near-zero scores suggest additivity.

### Cytokine Array and Analysis

Experiments were performed based on the protocol provided by bio-techne®. Briefly, 2 mL of Array Buffer 6 was added into each well of a 4-Well Multi-dish with membrane. After 1 hour incubation by rocking the membrane on the shaker, Buffer 6 was removed and the mixture of 1 mL sample (either supernatant from cell culture or cell lysate with indicated treatment) with 0.5 mL Buffer 4 was added into each well. Dish was covered and incubated overnight at 4 °C on the rocking shaker afterward. After incubation, each membrane was subsequently moved to individual plastic containers containing 20 mL of 1X Wash Buffer, and the dish was rinsed and dried thoroughly while three washes with 1X Wash Buffer for 10 minutes each on the rocking platform shaker was performed. After washing, the Detection Antibody Cocktail was prepared by adding 30 μL to 1.5 mL of 1X Array Buffer 4/6, and 1.5 mL was pipetted per well into the Multi-dish. The washed membranes were returned to the dish and covered and a 1 hour incubation at room temperature on the rocking shaker was conducted. Following this, the membranes were washed as previously described. Next, 2 mL of 1X Streptavidin-HRP was pipetted into each well, covered, and incubated for 30 minutes at room temperature. After further washing, a total of 1 mL of the prepared Chemi Reagent Mix was evenly pipetted onto each membrane, covered with the top sheet of the protector, and smoothed to remove air bubbles. After 1 minute incubation, signals were detected by ImageQuant™ LAS 4000 (GE healthcare). The intensity of each signal was quantified and recorded by ImageJ.

### TCGA analysis

Kaplan-Meier analysis and log-rank statistics were performed using R (version 4.2.0). Survival analysis is based on two grouping methods: High-PRDX6 and Low-PRDX6 groups, determined by the median expression of PRDX6, and samples with significantly upregulated PRDX6 and downregulated PRDX6 groups in tumors. Overall survival difference between two groups was evaluated using Kaplan-Meier analysis and verified by the log-rank test. Statistical significance was defined as a *p*-value < 0.05. The gene expression matrix and clinical information of the TCGA patients were downloaded from the UCSC Xena browser.

### Cell Growth Rate Assay

For each type of cell, 500 cells were seeded into 12 wells of each 96-well plate (7 plates total, 1 plate/day). After indicated days of incubation, plate of the day was washed by 100 μL D-PBS twice and incubated with 50 μL of 4% formaldehyde in D-PBS under room temperature, 100 rpm shaking for 10 minutes (Thermo, MaxQ 2000). After 10 minutes shaking, cells were washed again with 100 μL D-PBS and incubated with 50 μL of 0.05% crystal violet staining buffer (250 mL methanol, 750 mL MiliQ water, 0.5g crystal violet) under room temperature, 100 rpm shaking for 10 minutes. After shaking, cells were washed with DI water and air dried for 24 hours. Dried stained samples were resolubilized using 150 μL solubilization buffer (400 mL ethanol, 100 mL methanol, 100 mL MiliQ water) under 4°C, 100 rpm shaking for 1 hour. Resolubilized samples were further quantified by plate reader to measure the intensity of solubilized crystal staining (absorbance 530 nm, Molecular Devices SpectraMAX iD3).

### RNA Sequencing and Analysis

RNA was isolated from A223 cells treated for 16 hours using PureLink RNA Mini Kit (Thermo) following the manufacturer’s instructions. The concentration of each sample was determined using the QubitTM 3.0 Fluorometer (Thermo Fisher), and its integrity was assessed using an Agilent Bioanalyzer 2100 (Agilent Technologies). The Illumina TruSeq RNA Sample Preparation kit was utilized to create RNA sequencing libraries. The resulting library fragment lengths were confirmed using an Agilent Bioanalyzer 2100. Library quantification was performed using the QubitTM 3.0 Fluorometer, and sequencing was conducted at the Next-Generation Sequencing Facility at the University of Colorado BioFrontiers Institute using an Illumina HiSeq 2000 sequencing system. All sequencing libraries underwent multiplexing before sequencing. Reads were pruned using BBDUK from the BBMap suite v38.05 and mapped to hg38 human genome using HISAT2 v2.1.0. After mapping, alignment files were processed using SAMtools v1.3.1. FeatureCounts from the Subread software package v1.6.0 was used to count the total number of sequencing reads that aligned to each putative gene model. To determine which genes were differentially expressed, we used the R package DESeq version 1.34.1.

### *In vivo* mouse work and tumor injection

Tumor cells (1×10^5^ A223 or A223 PRDX6KD) were injected into wild-type (WT) C57BL/6 (B6) (Stock no. 000664, Jackson Laboratories). Both male and female mice (6-8 weeks) were used for the study. All mice were maintained under specific pathogen-free conditions in the vivarium facility of UPMC Hillman Cancer Center Animal Facility (Pittsburgh, PA). Animal work was approved by the Institutional Animal Care and Use Committee of University of Pittsburgh (Pittsburgh, PA).

Tumor cells were cultured in complete DMEM media supplemented with 10% fetal bovine serum (FBS), 20mM HEPES buffer, 1×antibiotic-antimycotic at 37°C CO_2_ incubator (5%) until 90% confluent. For tumor injection, cells were washed with phosphate buffered saline (PBS), trypsinized (0.01% Trypsin-EDTA, Fisher Scientific) and washed sequentially with complete DMEM media or PBS. Cells were suspended in PBS and 50% Matrigel Basement Membrane Matrix (Corning) to a final volume of 100µL and injected subcutaneously into one flank of each mouse. Tumor length and width were measured with calipers, and tumor volume was calculated as (length×width^2^)/2. Anti-PD-L1 (200µg/mouse/time, clone 10F.9G2, Leinco Technologies, Catalog#P371) diluted in PBS was given by intraperitoneal (i.p.) injection for 3 times (2-day interval). OKI-179 (60mg/kg/time) diluted in 0.1M citrate buffer was given by oral gavage for 9 times (2-day interval). To assess treatment effects, relative <u>c</u>hange in tumor <u>v</u>olume (RCTV) was calculated as the change in tumor volume (TV) from the start of treatment (TV_0_) to the TV at day n (the endpoint of control group) (TV_n_) divided by TV_0_ (RCTV=[TV_n_−TV_0_]/TV_0_). Based on the RCTV treated recipients were divided into R (RCTV<0), SP (0<RCTV≤1.5) and NR (RCTV>1.5). Recipient survival was monitored until mice reached experimental endpoints including severe tumor ulceration, tumor size >2cm in any dimension or other humane end points. Mice were euthanized in accordance with institutional guidelines.

### Graph Generation and Statistical Analysis

Both graph generation and statistical significance was created by BioRender. Data are represented as mean ± SD. Statistical analysis for data was performed as indicated and *p*-value < 0.05 was considered as statistically significant.

### Source Data Availability

Data supporting the findings of this study are reported in Supplementary Figures S1-S10 and Supplementary Tables S1-S2. All raw data corresponding to high-throughput approach (RNA-seq) are available through the NCBI’S Gene Expression Omnibus (GSE310671). All reagents and materials generated in this study are shown in Key Resources Table and will be available to the scientific community through Addgene and/or material transfer agreements. Further information and requests for resources and reagents should be directed to and will be fulfilled by the corresponding author: Xuedong Liu.

