## Supplemental Figures 1-9 for "PRDX6 Modulates Immune Checkpoint Inhibitor Response by Antagonizing Ferroptosis Induced By HDAC Inhibitors"

### Supplemental figure 1

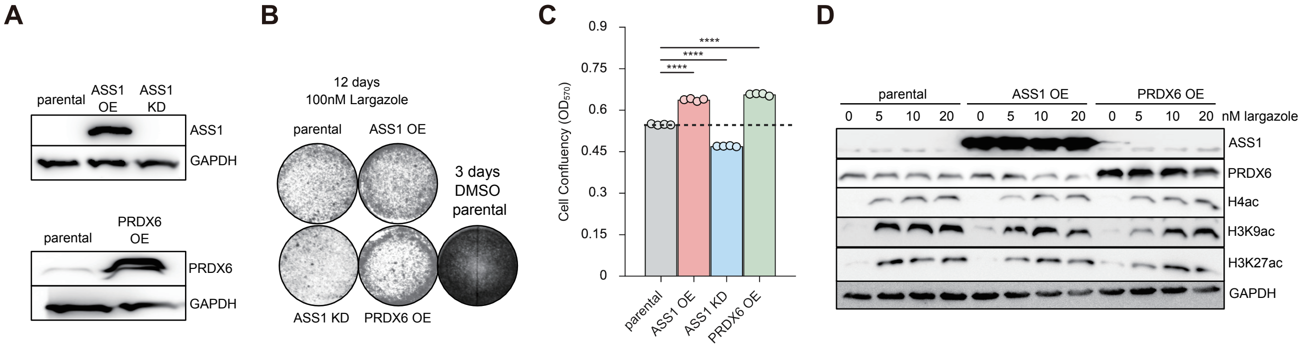

#### Supplemental figure 2

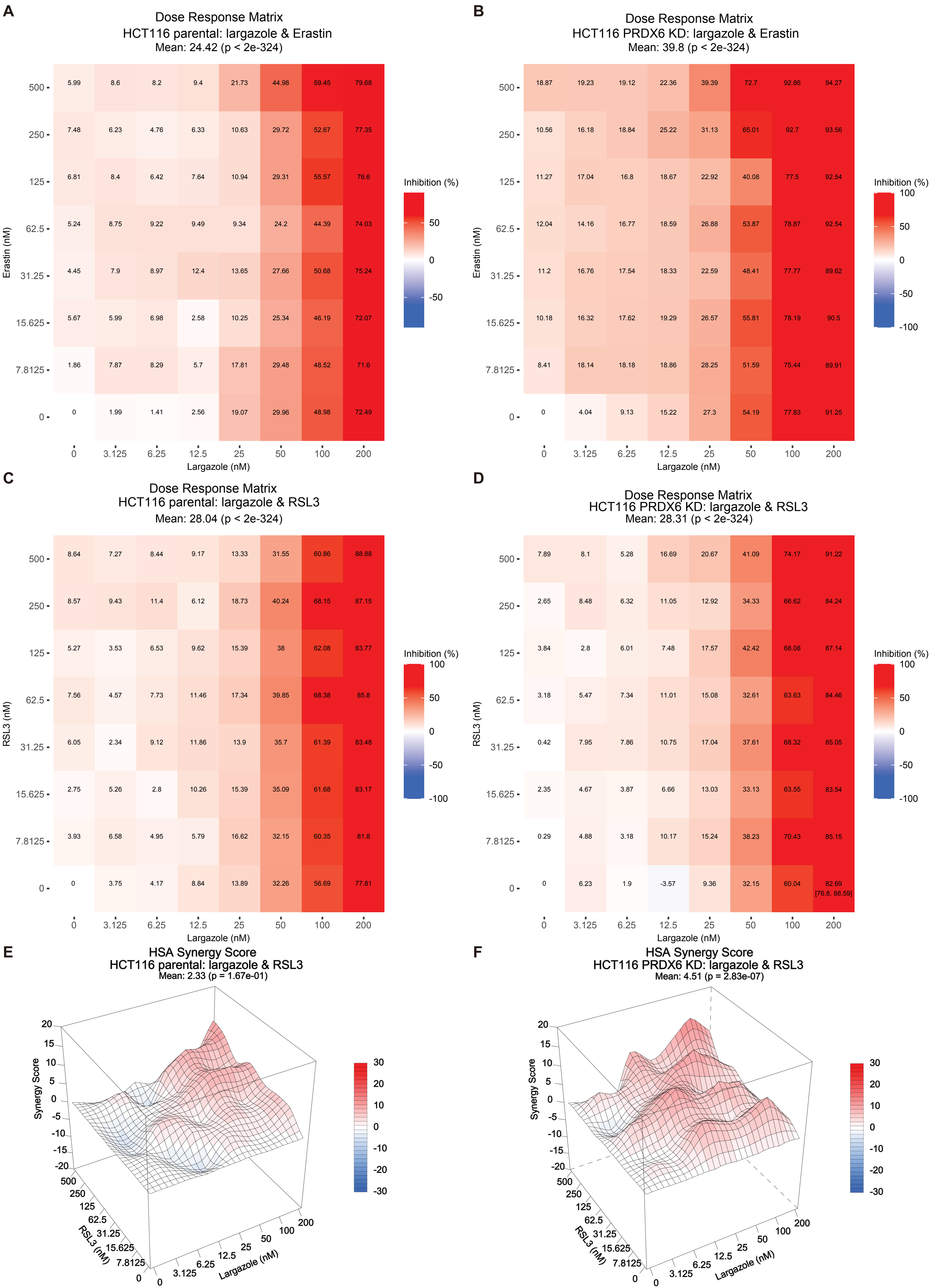

#### Supplemental figure 3

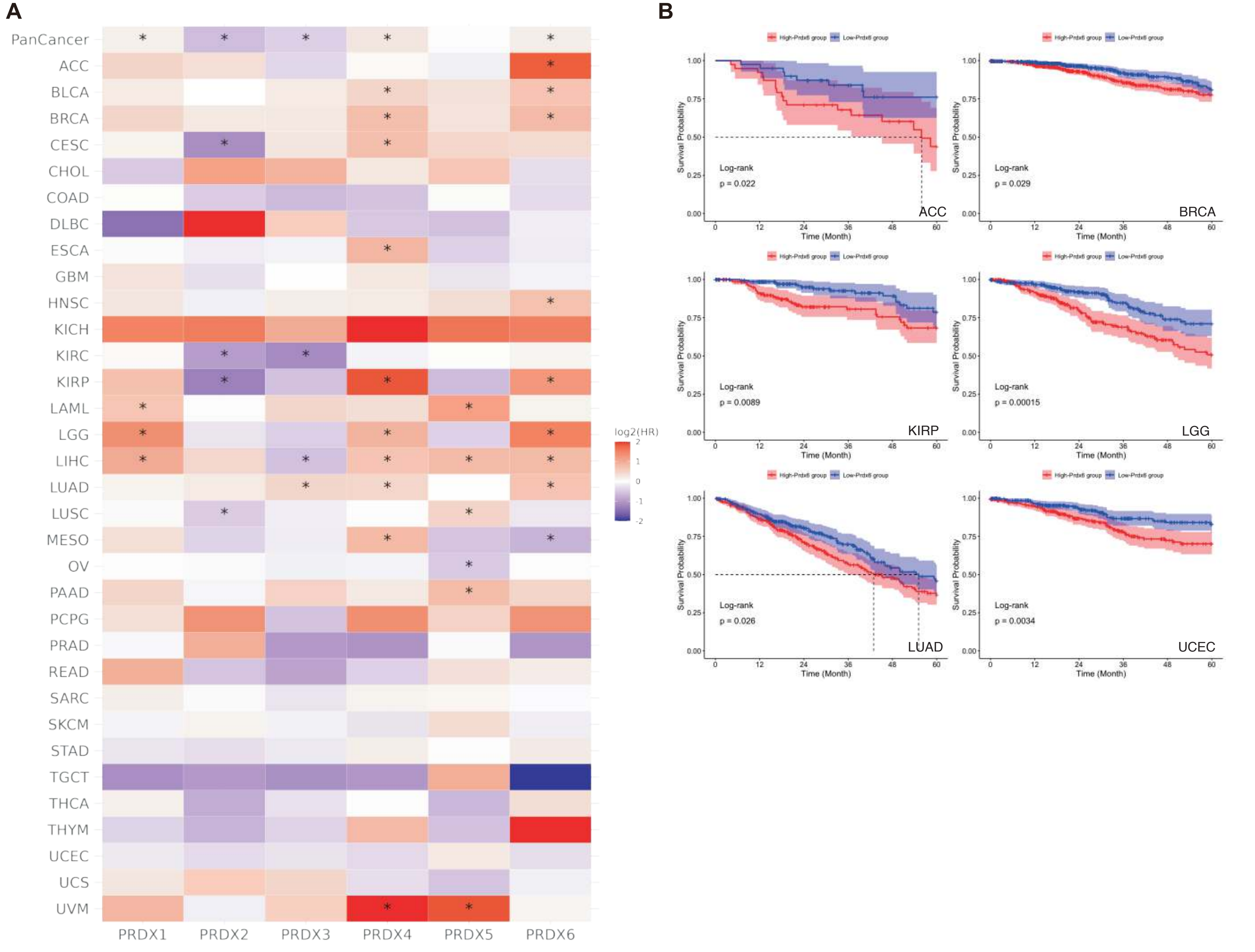

### Supplemental figure 4

A

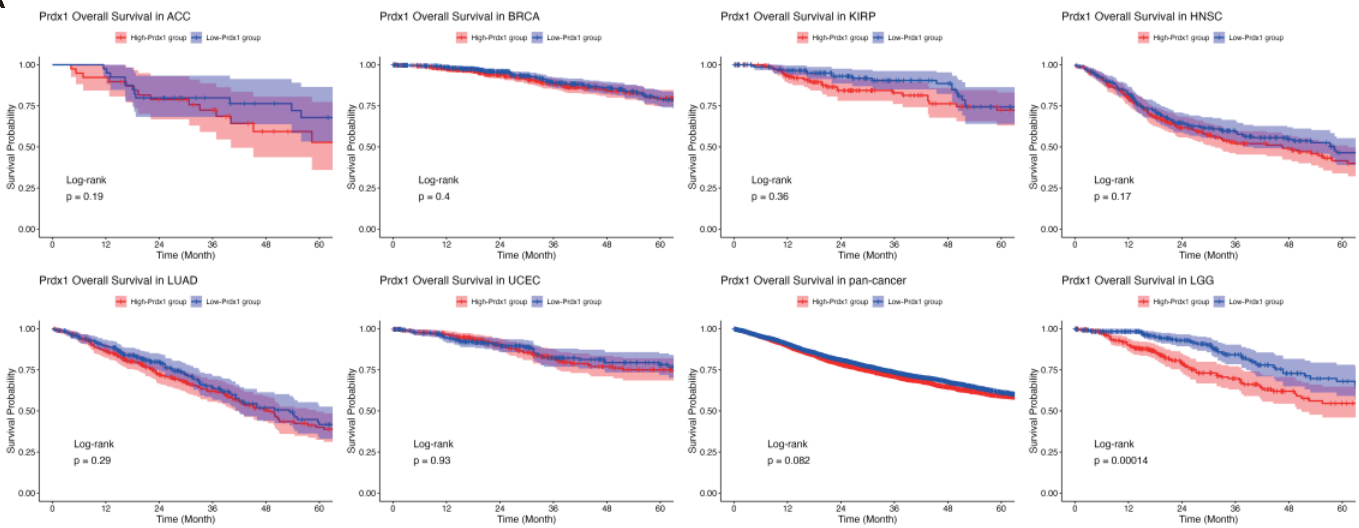


B

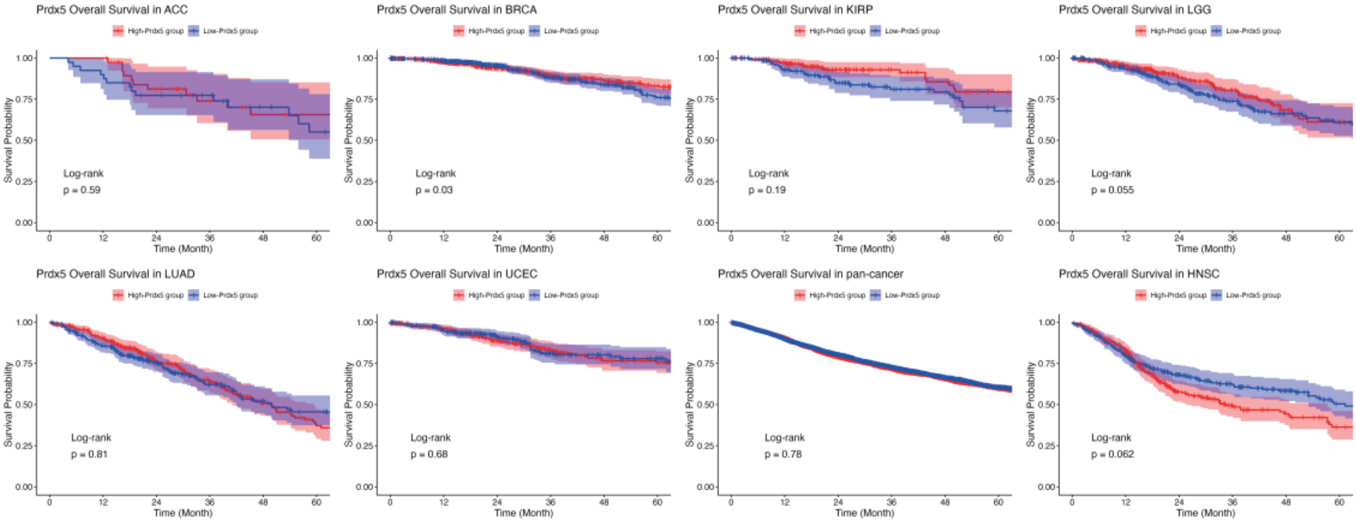

#### Supplemental figure 5

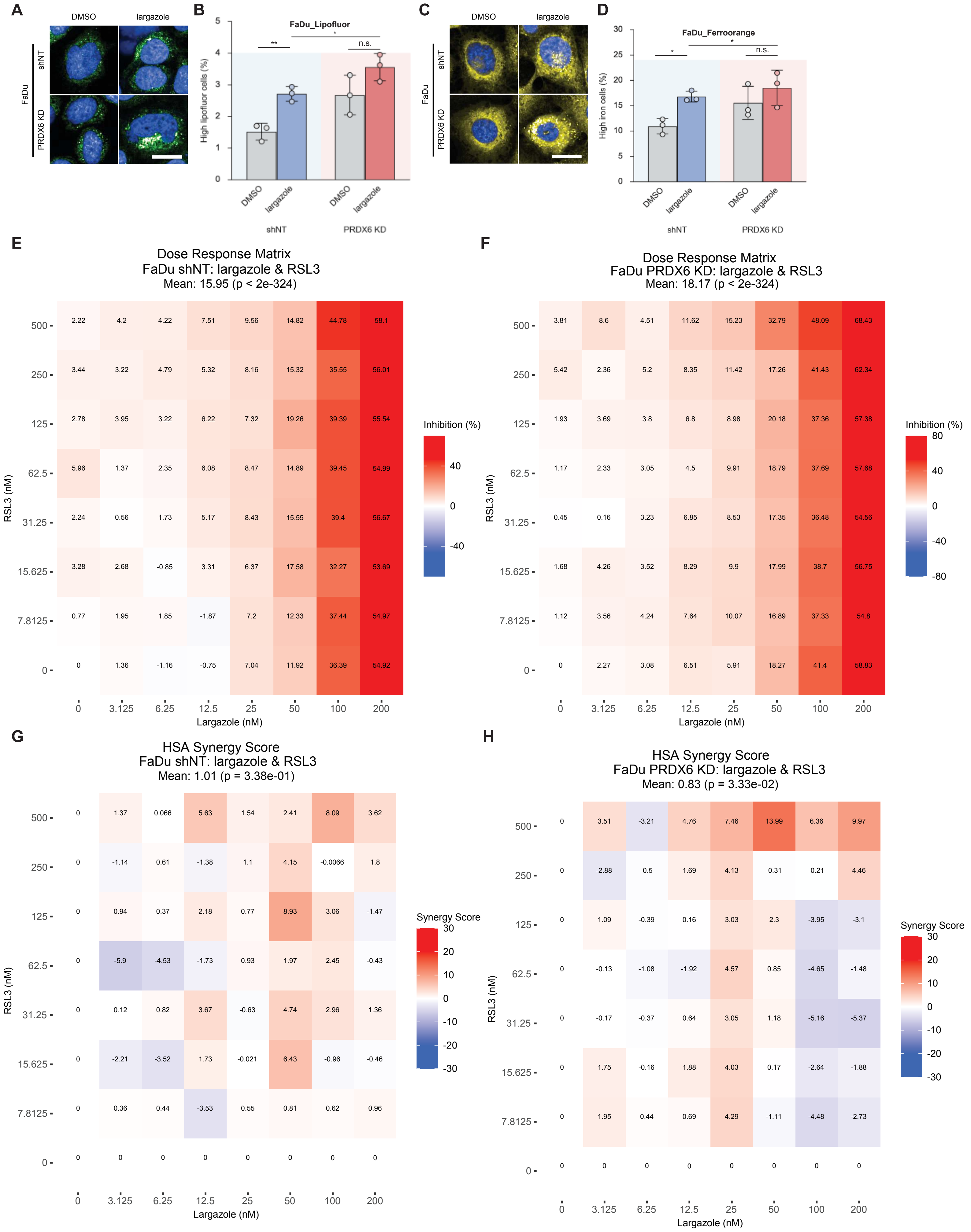

### Supplemental figure 6

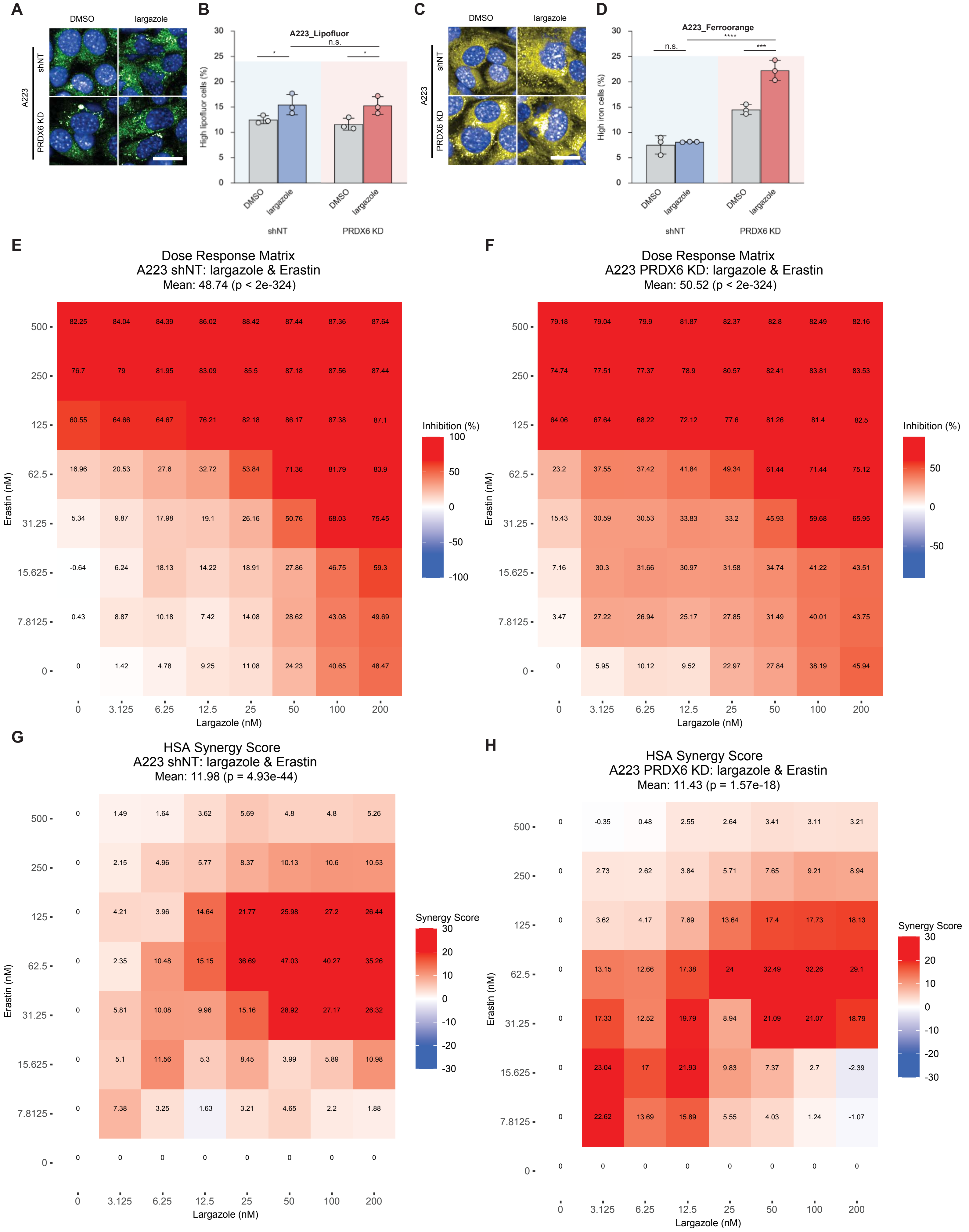

#### Supplemental figure 7

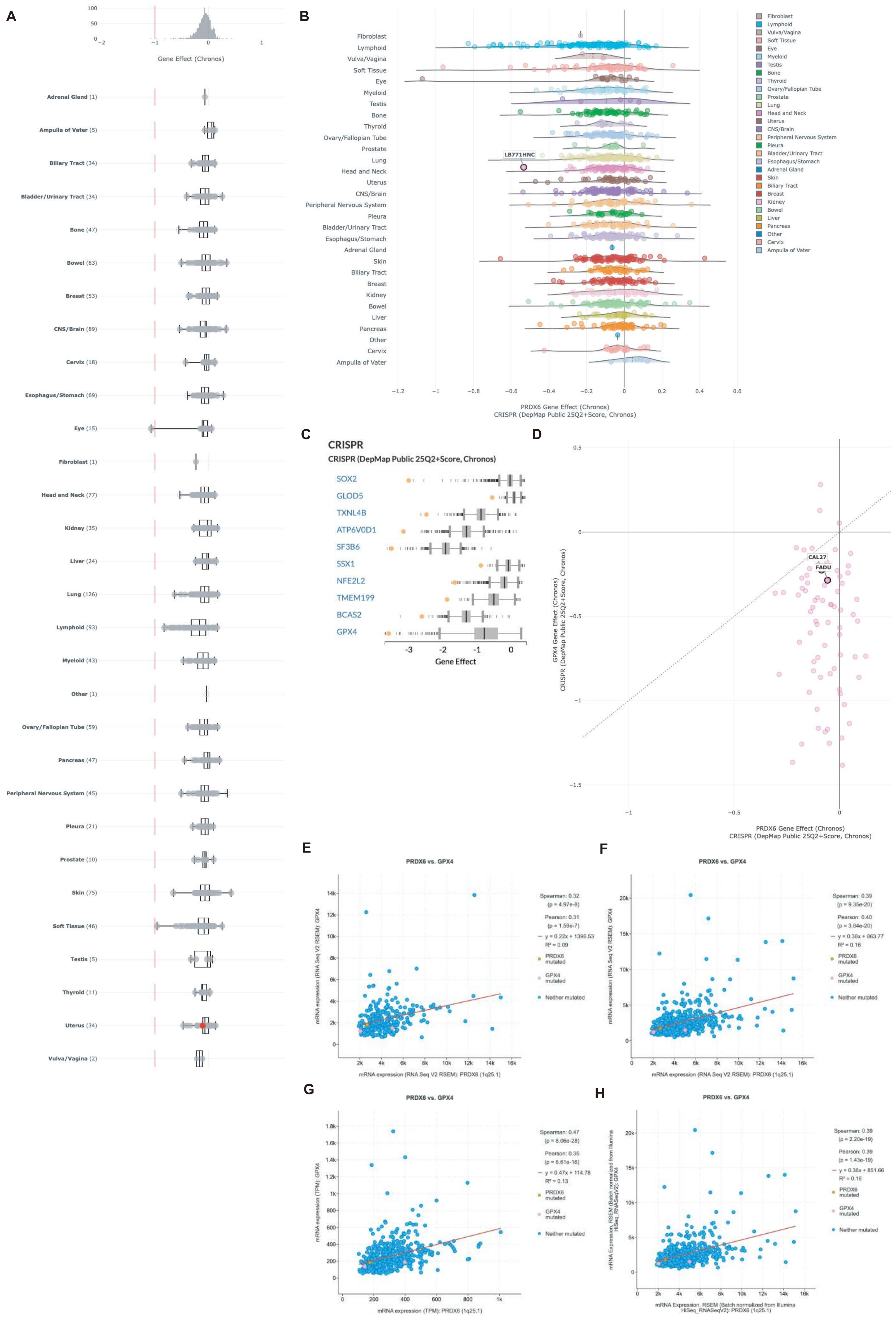

### Supplemental figure 8

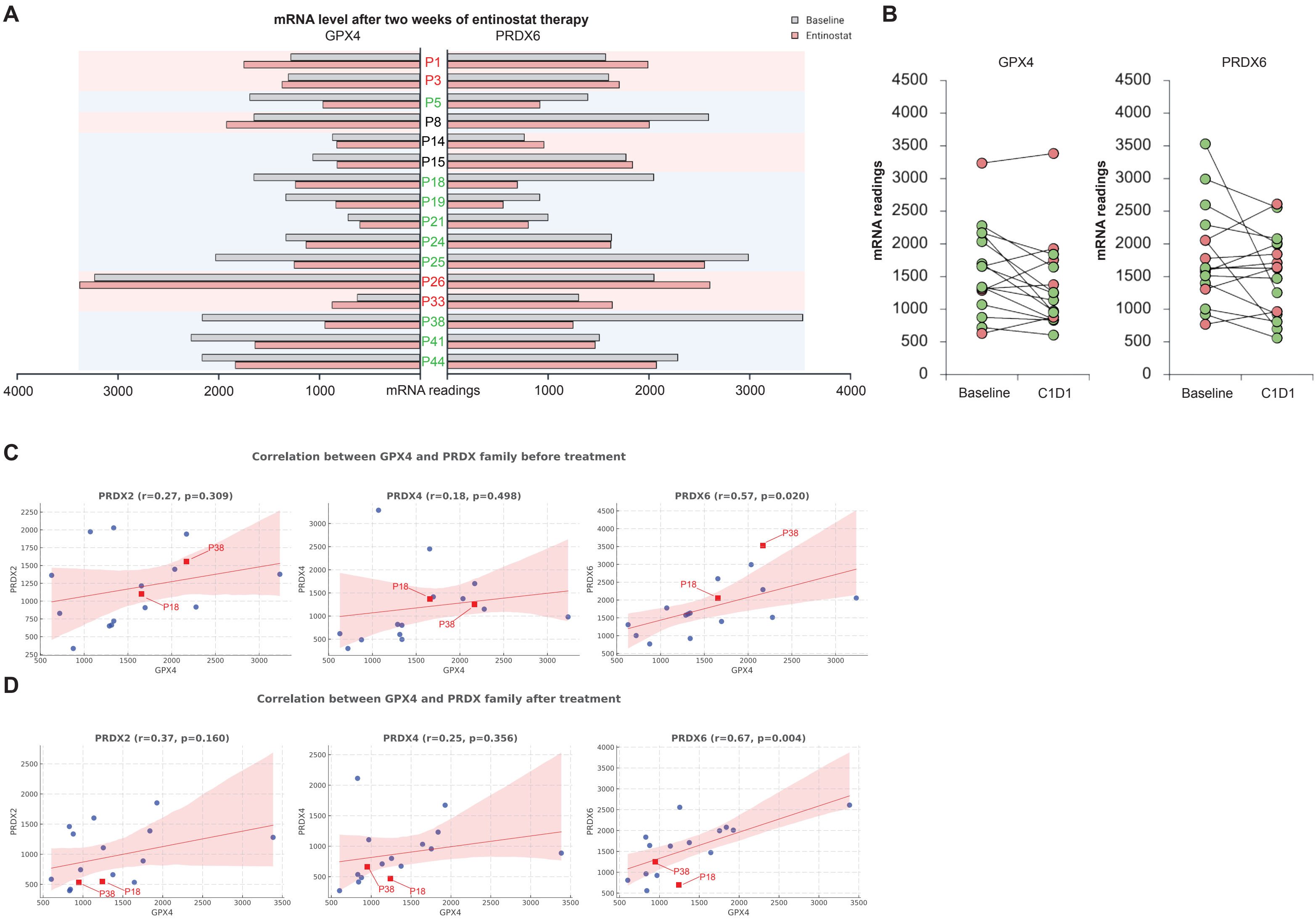

### Supplemental figure 9

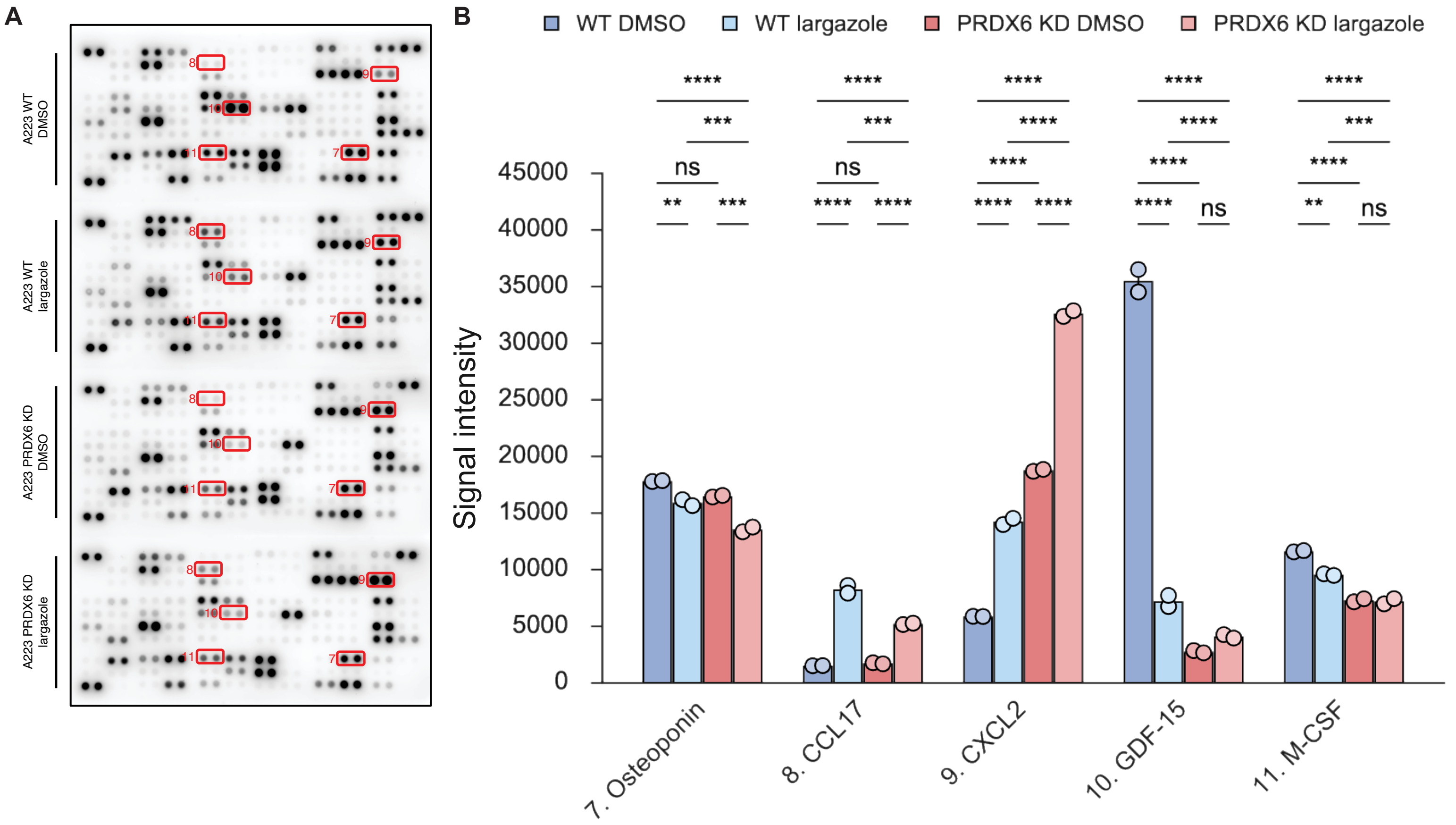
